# *Community* assesses differential cell communication using large multi-sample case-control scRNAseq datasets

**DOI:** 10.1101/2024.03.01.582941

**Authors:** Maria Solovey, Muhammet A. Celik, Felix R. Salcher, Mohmed Abdalfattah, Mostafa Ismail, Antonio Scialdone, Frank Ziemann, Maria Colomé-Tatché

## Abstract

Cell-cell communication is essential for physiological tissue function. In disease, this communication often gets disbalances by changes in the tissue cell type composition, fraction of cell engaged in communication and the rising or dropping expression levels of ligands, receptors and adhesion molecules. The changes in all these components of communication can be studied using single cell RNA-sequencing (scRNAseq) methods. With dropping sequencing costs, it is now possible to perform scRNAseq studies in larger cohorts of case and control samples to better address the heterogeneity of diseases. Here we present *community*, an R-based tool that is designed to perform differential communication analysis using scRNAseq between large cohorts of cases and controls. *Community* is able to reconstruct communication between different cell types both in the case and the control cohort of a dataset, and subsequently analyze which communication channels are affected in disease. *Community* is the first tool that integrates cell type abundance into the calculation of an interaction strength. *Community* is also able to disentangle the mechanisms underlying these changes, as well as detect interactions that are kept compensated by a sender or a receiver despite the disbalanced signaling from the counterpart. We tested *community* on two disease entities, ulcerative colitis and acute myeloid leukemia, using published scRNAseq datasets. We compared the performance of our tool to other differential communication pipelines, which *community* outperformed in speed and robustness. Overall, *community* is a fast, well-scalable, user-friendly R tool to assess differential cell-cell communication using large case-control scRNAseq datasets disentangling the driving mechanisms of communication shifts in disease.

## Introduction

Cell-cell communication is crucial for maintaining healthy physiological functions in tissues. In disease, the immune and non-immune niches, as well as the resident cell types, change their abundance and function, which results in an altered interaction among them^1–4^. Discovering these changes is important to better understand disease and identify new treatment strategies. Single cell RNA sequencing (scRNAseq) techniques allow to capture the shifts in the tissue composition, as well as in the expression of ligands, receptors and adhesion molecules within the individual cell types. ScRNAseq has become increasingly used in case-control types of studies where several cohorts with multiple samples are analyzed. This strategy allows better accounting for biological heterogeneity, which is especially relevant when working with human samples.

For the study of cell-cell communication changes between cases and controls, a differential communication analysis is needed. We previously published COMUNET, the first tool being able to perform differential communication analysis^5^. However, COMUNET performs differential communication analysis of one control versus one case sample, while often many samples are analyzed in cohorts. An algorithm that is able to scale to multiple samples easily is thus needed. Several other cell-cell interaction tools are able to perform differential analysis^6–9^. The major limitation of these tools is that the disease-driven shifts in the cell type abundance are not considered while analyzing differential communication. These shifts can though be massive and can vastly affect the overall communication dynamics within the diseased tissue. To maintain homeostasis and proper tissue function, several mechanisms adapt at the cellular level. First, cells (especially immune cells) can infiltrate a tissue, resulting in an increase of their cell type abundance. Second, cell types can decrease in their abundance due to e.g. damage, aging or clearance via the immune system. The remaining cells can adapt their ligand and receptor expression levels in an attempt to keep the physiological levels of interaction stable and compensate for the shifts in the tissue composition. It is of great importance to take these processes into consideration in the cell-cell communication analysis.

Here we present *community*, an R package that performs differential communication analysis in large scRNAseq case-control studies. *Community* is a fast and user-friendly tool which is not limited by the number of samples in the case and control cohorts that can be processed. *Community* does not rely on prior differential expression analysis, but captures changes in cell type abundance, fraction of active cells, and ligand-receptor and adhesion molecules expression. *Community* can thus recognize not only interactions that are differential between cases and controls, but also distinguish between unchanged interactions and those that are being compensated through the interaction of the several components.

We exemplify the differential communication analysis applying *community* to two very distinct disease entities: ulcerative colitis (UC) and acute myeloid leukemia (AML). We show that UC undergoes an upregulation in communication intensity, including both epithelial, vascular and immune compartments, which is in line with the physiology of the disease manifesting increased inflammation. In contrast, on the AML datasets we show a decrease in communication activity, especially within the immune compartment. This observation is in line with the described effect of immune silencing via the cancer cells. *Community* identifies differential interactions even in very small populations of cells, and it can as well detect complex interactions. We compare *community* to other two broadly used communication algorithms: CellPhoneDB^6^ and NicheNet^10^, using their differential communication pipelines and show that *community* is faster, more robust to outlier signals and is able to capture hidden compensatory effects.

The intuitive output of *community* makes it possible to track shifts in communication among cell types in diseased tissues compared to controls, visualize and investigate individual top differential interactions, as well as identify the driving components underlying the communication changes. The large-scale studies of shifts in cellular interactions will help understand biological mechanisms driving disease progression at a new level of complexity.

## Results

### Theoretical basis

To assess communication between case and control samples using scRNAseq, *community* uses ligand-receptor and adhesion molecule interactions between cell types in a tissue (Fig. 1a). It works with a flexible build-in database of known ligand-receptor and adhesion molecule interactions adapted from *OmniPath^11^*, which can easily be expanded by the user (Supplementary Fig. 1). To construct the database, we utilize two main interaction databases from *OmniPath*: ‘ligrec extra’ and ‘curated ligand-receptor interactions’.^11^ We break down all complex interactions into their individual components, such that all interacting pairs consist of only one ligand and one receptor. The components of adhesion molecule pairs or other non-directional interactions are randomly labeled as “ligand” or “receptor”, but this directionality can be ignored in these cases. The generated database comprises 6,941 entries, of which 4,157 are ligand-receptor pairs, while 2,784 are between adhesion molecules. Each ligand, receptor and adhesion molecule in the database has a functional annotation obtained from *OmniPath*.

**Fig. 1.**
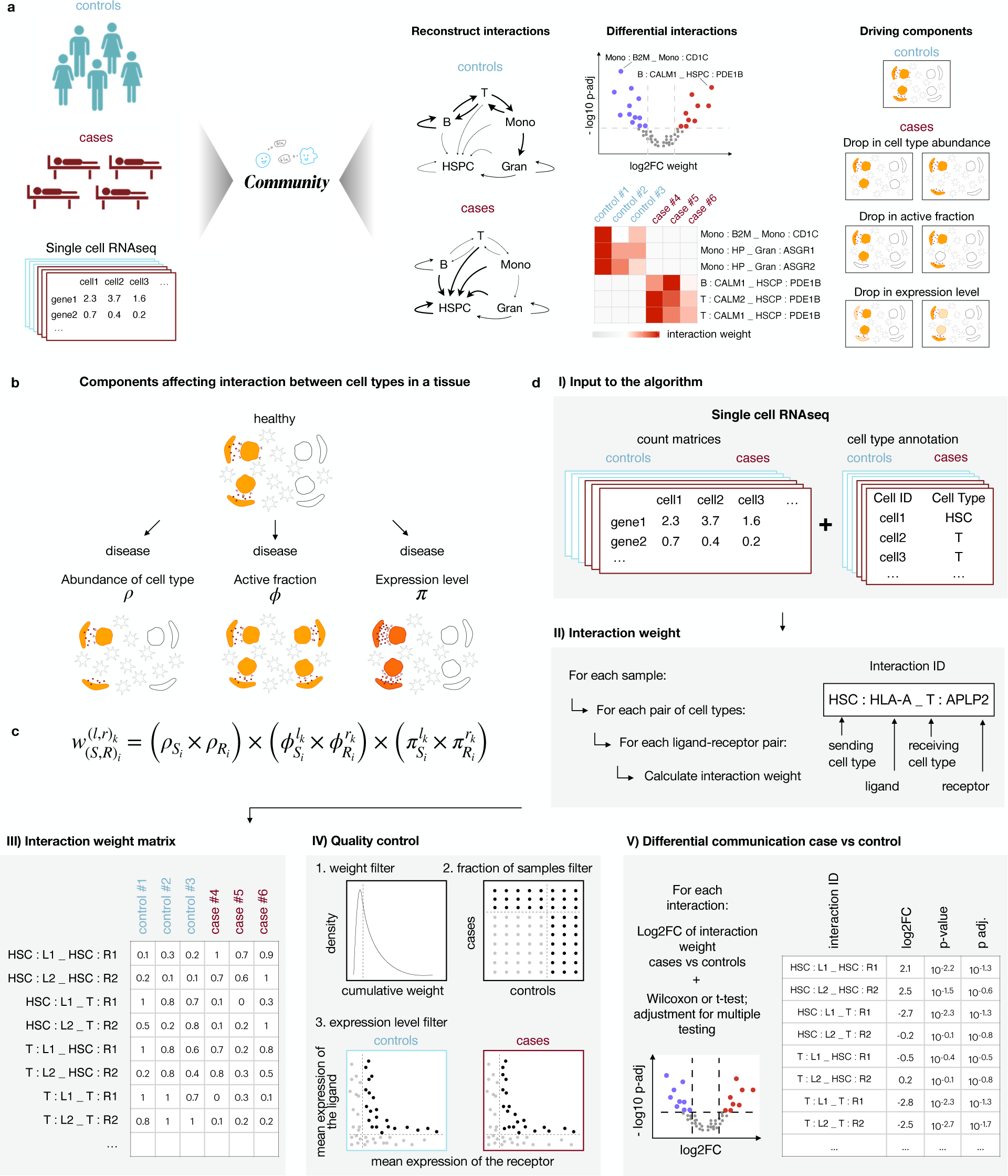
Overview of the *community* tool. **a**, *Community* overview. **b**, Components affecting interaction between cell types in a tissue. **c**, Formula to calculate an individual interaction. **d**, *Community* workflow.

*Community* is unique in its definition of the communication weights, which are defined by three major contributors to the communication intensity: cell type abundance *ρ*, active fraction *ϕ*, and mean expression level within the active fraction *π* (Fig. 1b). Cell type abundance is frequently affected in disease due to shifts in the cell type composition. If one or both communicating cell types are shrinking or expanding, this will affect the tissue-level intensity of the interaction between these cell types. *Community* takes this into account by including changes in cell type abundance *ρ* in the differential communication analysis. The active fraction *ϕ* is defined as the fraction of cells in a particular cell type that has a non-zero expression of the ligand or the receptor of interest. In a disease, if the active fraction of a cell type expands or shrinks, the communication with this cell type may get affected, even if the cell type abundance and the expression level of the individual cells remain unchanged. Finally, the mean expression level *π* of ligands and receptors within the active fraction reflects the intensity of the interaction, and it is the most intuitive and well-studied component that affects communication intensity. All communication tools designed to date calculate communication strength based only on average expression per cell type (pseudobulk), which is a product of *ϕ* and *π*. However, these pseudobulk expression changes do not capture the disease-driven change in the cell type abundance, nor give the specific information whether the alteration in the psedobulk-level expression was due to the change in the fraction of cells expressing a gene or in the expression levels themselves.

Based on these three components, the weight of an interaction between two cell types (sending S and receiving R), mediated by a particular ligand-receptor pair (*l*, *r*)_*k*_ (or adhesion molecule pair (*l*, *r*)_*k*_) in a particular sample is defined as the product of the three components:

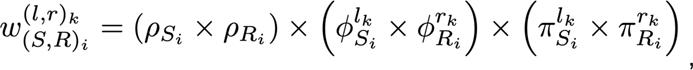

*ρ_Si_* is the relative abundance of the sending cell type, *ρ_Ri_* is the relative abundance of the receiving cell type, *ϕ^lk^_Si_* and *ϕ^rk^_Ri_* are the relative active fraction of the ligand *l_k_* in the sending cell type and the receptor *k* in the receiving cell type respectively, and *π^lk^_Si_* and *π^rk^_Ri_* are the relative mean expression of the ligand *l_k_* in the active fraction of the sending cell type and the receptor *k* in the active fraction of the receiving cell type respectively (Fig. 1c; Supplementary Fig. 2, Supplemental Methods). The interaction weight *ω^lrk^_SRi_* can have a value between 0 and 1.

*Community* is designed to work with scRNAseq data from case-control experiments. Its input is a pre-processed (i.e. normalized and batch corrected, if needed) scRNAseq dataset with cell types already annotated (Fig. 1d Input to the algorithm; Supplementary Fig. 2). Like in all cell-cell interaction analyses, cell types have to be present in both cases and controls, thus merging fine-grained cell type annotations might be needed. For diseases such as cancer, the malignant cell types should be annotated to the cell type category of their healthy counterparts.

In the first step of the algorithm, for each sample, we calculate the interaction weight between each pair of cell types for each ligand-receptor pair (LRP) from the database (Fig. 1d Interaction weight; Supplementary Fig. 2). Three thresholds are applied: a cell type is considered as captured in a particular sample if there is a minimum number of cells in that cell type; this cell type is considered to have a big enough active fraction if there are a minimum number of cells in that cell type expressing a particular ligand or receptor; a cell is considered as active if it reaches a minimum expression level of the ligand or receptor. Every interaction gets an ID constructed as *sending cell type : ligand _ receiving cell type : receptor*. This step is repeated for all samples, which results in an interaction matrix with sample IDs in the columns and interaction IDs in the rows (Fig. 1d Interaction weight matrix).

In the quality control (QC) step, the quality of the interactions is evaluated to filter interactions that are either too weak or are not present in a sufficient amount of samples (Fig. 1d Quality control). For this, we apply three filters: cumulative interaction weight over all samples, fraction of samples in which an interaction is present, and mean expression of the ligand and receptor per cohort (see Methods section).

In the final step of our analysis, we calculate differential interactions between the case and the control samples (Fig. 1d Differential communication case vs control). For each interaction, we first calculate the log2 fold change between cases and controls and the respective p-value. To calculate the p-value, we use either the t-test^12^ on the log2 transformed weight values, recommended for smaller cohorts, or the Wilcoxon test^13^ on the actual weight values, recommended for bigger cohorts. We correct for multiple testing using the FDR method^14^. After this, significantly differential interactions can be visualized and further studied.

The output of *community* contains, most importantly, the interaction weight matrix and the interaction annotation matrix with information on the individual components used for the weight calculation, QC metrics, as well as statistics for differential communication analysis. *Community* also implements intuitive options for visualization, such as for example vulcano plot of differential interactions, heatmap of top up- and downregulated interactions, and forest plots for analysis of changes in the individual components (Fig. 1a). The versatile output of *community* makes it possible to easily navigate from the top-level analysis of global changes in cellular communication between cases and controls, down to the detailed investigation of individual interactions, allowing to identify the components driving the change.

### Changes in communication in ulcerative colitis vs healthy colon

To demonstrate the capabilities of *community*, we applied it to a previously published scRNAseq dataset of primary human ulcerative colitis (UC) patients and healthy controls^15^ (Fig. 2a, Supplementary Fig. 3a-d). After filtering the dataset contained 17 UC and 11 healthy control samples, as well as 13 cell types (Fig. 2a, Supplementary Fig. 3a-d).

**Fig. 2.**
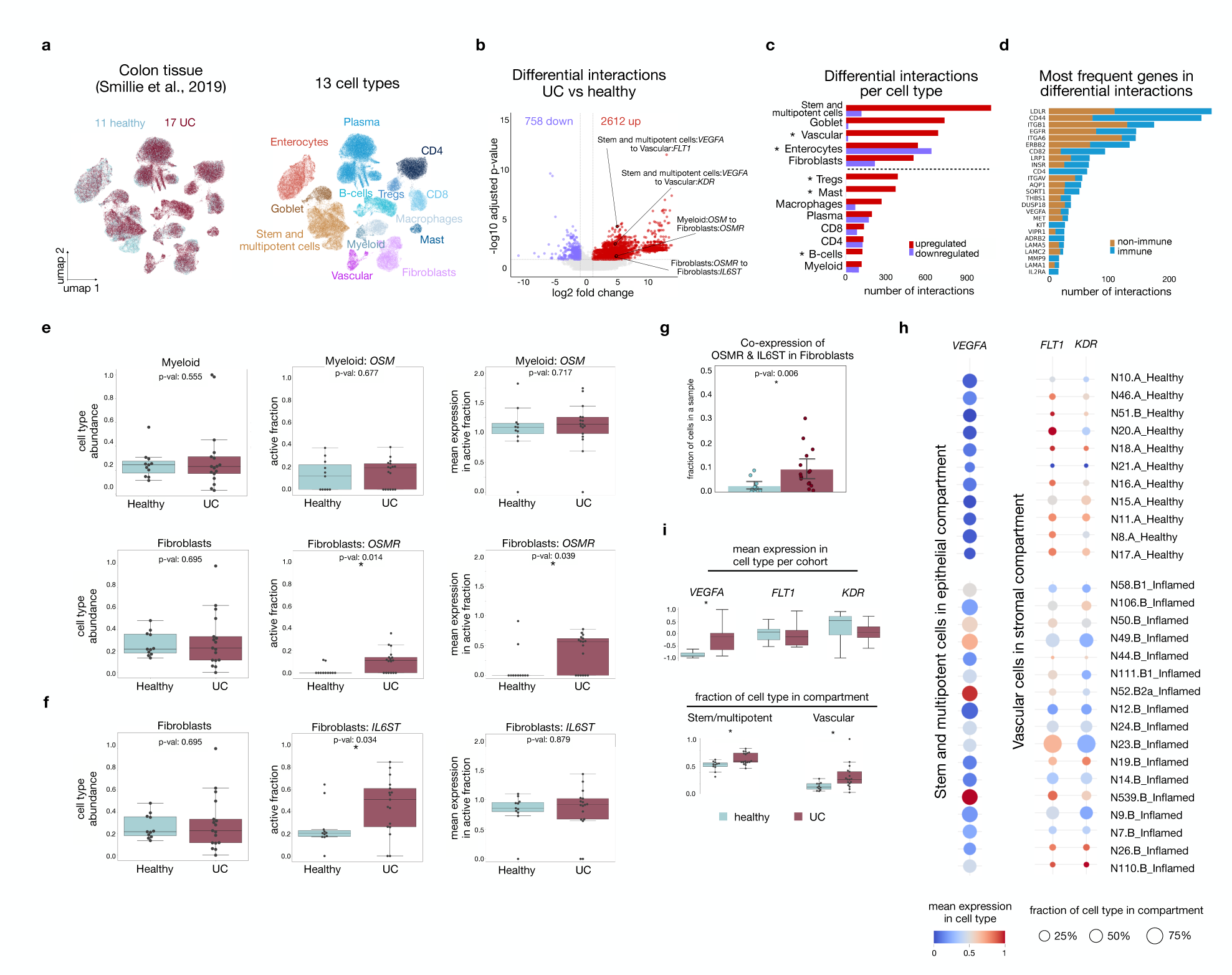
Communication in inflamed ulcerative colitis (UC) and healthy colon tissue. **a**, UMAPs colored by health status (left panel) and cell types (right panel). **b**, Volcano plot of differential interactions with highlighted relevant interactions. **c**, Number of differential interactions per cell type. Asterisk makes cell types with altered cell type abundance. **d**, Ranking of most frequently occurring genes in significant interactions. The box-lots in **e** and **f** present the cell type abundance, active fraction and mean expression in active fraction for relevant genes in the selected cell types. P-values were calculated using t-tests (p-val. <0.05), with an asterisk marking significance. **g**, Co-expression of OSMR and IL6ST over all Fibroblasts. The p-value was calculated using t-tests, with the asterisk marking significance (p-val. <0.05). **h,** Dot-plot of *VEGFA* expression in stem and multipoint cells and *FLT1* and *KDR* expression in the vascular cells. The color represents mean gene expression for a cell type per sample. Dot-size represents the fraction of the respective cell type in the cell-specific compartment. **i** Box-plots for the mean expression and fraction of cll type for the same genes in same cell types as **h.** Statistics were done by performing t-tests. Significant findings (p-val. <0.05) are labeled with an asterisk.

Using *community,* we identified 15,675 interactions that passed quality control, with 2,612 being significantly upregulated and 758 downregulated in UC (Fig. 2b; Supplementary Fig. 3e). In UC, as an inflammatory bowel disease, we expect to observe overall increased levels of communication, due to influx of immune cells and inflammatory cytokine production. Our analysis revealed a predominance of non-immune cell types in upregulated interactions, specifically stem and multipotent cells, followed by goblet cells, vascular cells, enterocytes and fibroblasts (Fig. 2c; Supplementary Fig. 4a). Of those, for stem and multipotent cells, goblet cells, and fibroblasts the change in communication was driven by an increase in active fraction and mean expression, whereas for enterocytes and vascular cells the shift was additionally driven by a strong change in cell type abundance (Supplementary Fig. 5). This is consistent with the vascularization of the inflamed tissue and the loss of enterocytes.^16,17^ Among immune cell types, T-regs and mast cells showed the highest number of upregulated interactions (Fig. 2c). These were driven by all three components (Supplementary Fig. 5), consistent with an immune activation. For the rest of the immune cell types, changes in communication were not caused by cell type abundance, and only infrequently driven by changes in active fraction or mean expression (Supplementary Fig. 5). Instead, the changes were driven by the interacting counterparts of those cell types (Supplementary Fig 5).

Next, we explored individual ligands and receptors in the differential interactions (Fig. 2d), identifying *LDLR*, *CD44*, *ITGB1* and *EGFR*, which are known to play critical roles in UC and other related pathologies of the intestine by driving inflammation.^18–21^ Most of the top ligands and receptors were used by both non-immune and the immune compartments. Others, like *CD4* or *KIT*, were immune compartment specific (Fig. 2d). We observed that for different cell types, individual ligands and receptors can contribute to the change in the interaction strength via different components. For example, *CD44* is used as a receptor by 11 cell types (Supplementary Fig. 4b). Its contribution to the increased interactions in the regulatory T cells, mast cells and B cells is driven solely by an increased cell type abundance of these cell types. In contrast, goblet cells, macrophages and stem and multipotent cells increased their active fraction expressing *CD44*. For the other cell types, all three components were predominantly stable and the change in interactions was driven by the sending counterpart cell types (Supplementary Fig. 4b). These results illustrate how various shifts in communication are regulated, and that it is important to consider all aspects of cell-cell interaction to gain biologically relevant analysis.

As proof of concept, we confirmed the upregulated interaction *Myeloid:OSM_Fibroblasts:OSMR* between myeloid cells expressing Oncostatin M (*OSM*) and fibroblasts expressing the Oncostatin M receptor (*OSMR)* (Fig. 2b,e,f, p.adj = 0.044, log2FC = 4.798). This aligns with the findings of the original publication, which have been confirmed with confocal microscopy^15^. *Community* identified that the overall increase in communication was driven by the expansion of an *OSMR*-expressing subpopulation of fibroblasts, as the fraction of fibroblasts expressing *OSMR* increased from 0% (healthy) to 17% (UC) (Fig. 2e). Numerous publications show that OSMR activation is accompanied by a co-activation of its heterodimeric counterpart IL6ST^22^. To test if this is the case, we compared co-expression levels of these two genes and observed a higher fraction of co-expression in fibroblasts in UC compared to healthy controls (p.val = 0.006) (Fig. 2g; Supplementary Fig 4c). Also, the *community* database marks the *OSMR-IL6ST* pair as a complex, and *community* captured the *Fibroblasts:OSMR_Fibroblasts:IL6ST* interaction as significantly upregulated (p.adj = 0.049, log2FC = 4.747), which is likely a consequence of the co-expression (Fig. 2f,g; Supplementary Fig 4c).

Besides exploring overall changes in communication and identifying components driving shifts in interactions already described in the initial publication, we also discovered novel interactions. Among them, we found a significantly upregulated interactions “*stem and multipotent cells:VEGFA_vascular:FLT1”* a n d “ *s t e m a n d m u l t i p o t e n t cells:VEGFA_vascular:KDR”* (Fig. 2b). *Community* identified these interactions to be driven by the increased active fraction and mean expression of *VEGFA* in stem and multipotent cells, as well as the increased cell type abundance of vascular cells (Supplementary Fig 4d). *VEGFA* is a known driver of vascular growth and pathological angiogenesis, which is often overexpressed in response to inflammatory hypoxia^16^. Our analysis shows increased mean expression of the PHD family (HRE complex members) which are the upstream pathway of *VEGFA*, in the stem and multipotent cells in the UC cohort. (Supplementary Fig 4e). As this interaction is driven by the abundance of vascular cells it was not captured in the original publication by Smillie et al.

These results showcase that *community* can be applied to case-control data sets of inflammatory disease. It allows for the identification of known interactions and enables the user to gain a deeper understanding of shifts in interactions. It is also capable of identifying complex interactions such as OSM-OSMR/IL6ST. Additionally, it can detect previously missed interactions that provide insights into disease specific communication patterns, which could aid in identifying novel therapeutic targets

### Changes in communication in AML vs healthy bone marrow

To assess the versatility of *community*, we also applied it to a cancer case-control data set, where by nature malignant cells are not present in the healthy tissue. This published human bone marrow scRNAseq dataset comprised 7 individuals with acute myeloid leukemia (AML) at diagnosis and 6 healthy controls^23^. We defined 8 cell types: hematopoietic stem and progenitor cells (HSPC), monocytes, granulocytes, dendritic cells (DC), erythrocytes, B-cells, T-cells and natural killer cells (NK) (Fig. 3a; Supplementary Fig. 6a-d).

**Fig. 3.**
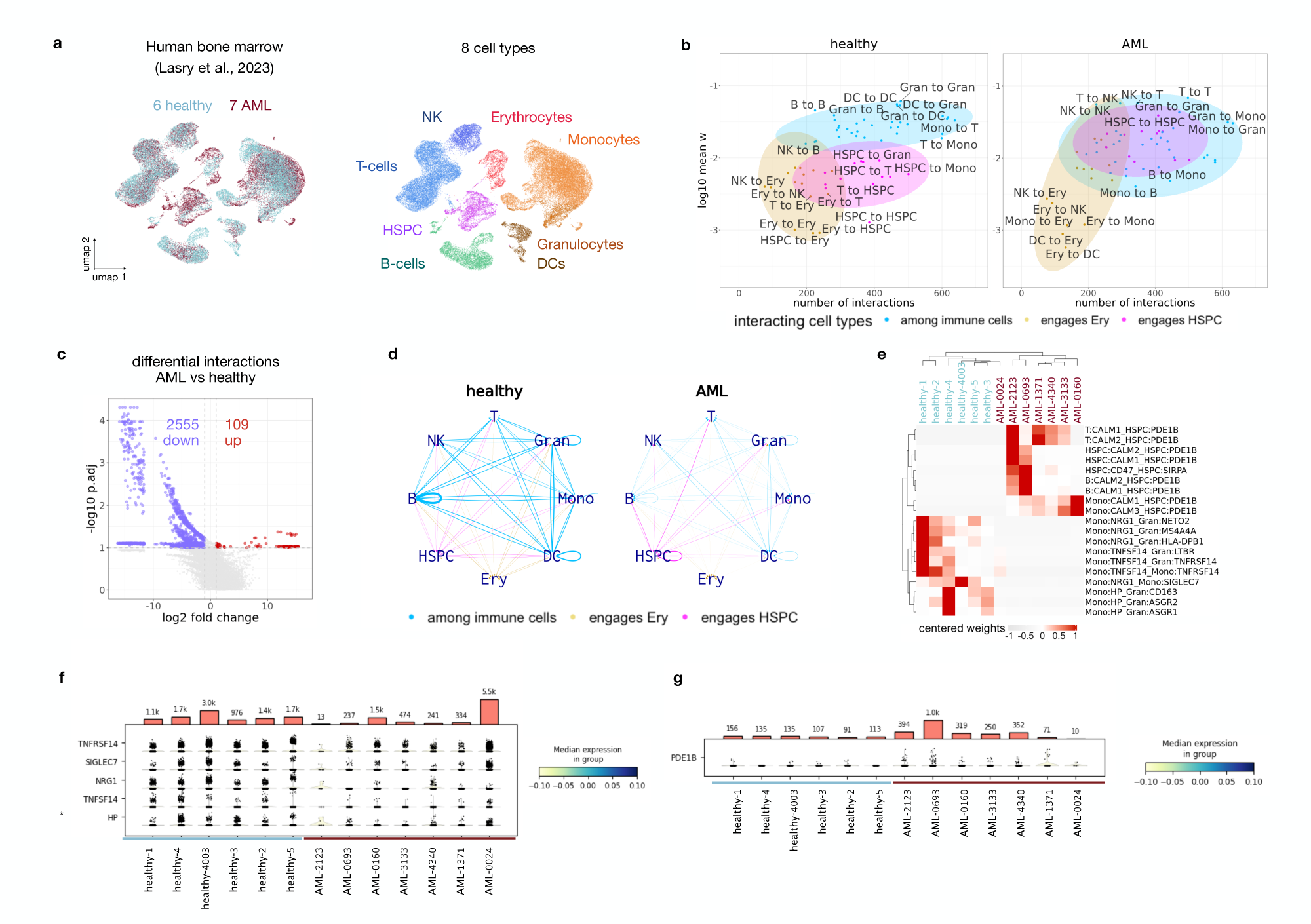
Communication in healthy and AML bone marrow. **a**, UMAP of the human bone marrow scRNAseq dataset coloured by health condition (left panel) and cell type (right panel). **b**, Good quality interactions groups by cell type for healthy (left panel) and AML (right panel) cohort. **c**, Volcano plot of differential interactions (AML vs healthy). **d**, Network graph of differential interactions of healthy (left panel) and AML (right panel) cohort. One line represents cumulative weight of all differential interactions between the two connected cell types **e**, Heatmap of top up-and down-regulated interactions. **f**, Stacked-violin plots of the ligands engaged in the top down-regulated interactions of the panel **e**. **g**, Stacked-violin plot of the receptor engaged in the top up-regulated interactions of the panel **e**.

After running *community* and filtering out interactions that did not pass quality control, we identified 22,645 good quality interactions. First, to get a general impression of the communication results, we grouped the interactions per cell type. To do that, we aggregated the interaction weights of all communication edges between every pair of cell types, and then summarized them into three broad interaction categories: “engaging HSPC”, “among immune cell types”, and “engaging erythrocytes” (Fig. 3b). The engaging HSPC category included classes that had HSPC either as a sending or as a receiving cell type, or both. The category “among immune cell types” included classes where both sending and receiving cell types were immune cells. The “engaging erythrocytes” category had erythrocytes either as a sending or as a receiving cell type or both. In the healthy cohort, we observed that the class of interactions in the “among immune cell types” category had higher average weights compared to the AML, whereas classes engaging HSPC or erythrocytes tended to have lower weights. In the AML cohort instead we observed a drop in the average weights of immune interactions, indicating less immune-driven communication in these patients (Fig. 3b). In AML, the HSPC-engaging classes had higher interaction weights compared to the healthy control (Fig. 3b). These results align with known biology, as the immune compartment is most represented in healthy BM, therefore we expect it to contribute most to the overall communication in the tissue. In AML though, due to the differentiation block^24^ and immune suppression^25^, we expect to observe silencing of the immune-driven communication. Additionally, due to the expansion of the blasts and their reported suppressive interaction activity towards other compartments, we expect to observe increased interactions engaging HSPCs in AML.

We next proceeded to the differential communication analysis. We identified 2,664 differential interactions, 109 upregulated in AML and 2,555 downregulated (Fig. 3c). The majority of the downregulated interactions involved interactions of immune cell types among each other, while the upregulated interactions predominantly involved interactions between HSPCs and immune cell types (Fig. 3d). Like many cancers, leukemic cells avoid immune destruction as a cancer hallmark, by impairing immune response due to several mechanisms (e.g. T-cell exhaustion and HLA-loss), therefore it was to be expected to see an overall decrease in communication, especially involving immune cells.^25^,26 To further study mechanisms of decreased immune communication we checked which components are most prominent in each cell type. In this dataset, DCs most frequently showed drop in all three components: cell type abundance, active fraction and the mean expression (Supplemental Fig. 7). HSPC and erythrocyte precursors showed upregulated cell type abundance and downregulated active fraction, while B-cells, monocytes and granulocytes showed changes in active fraction and occasionally in mean expression. The T- and NK-cells showed only occasional changes in active fraction and mean expression (Supplemental Fig. 7). Among the top downregulated interactions, we found those engaging monocytes expressing *TNFSF14*, *NRG1*, *HP, TNFRSF14, and SIGLEC7*, predominantly driven by the reduction in the active fraction of these genes in the monocytes. (Fig. 3e,f; Supplementary Fig. 6f).

These genes are immune activating, and loss of the monocytes expressing these genes is an indication of an immunosuppressive milieu within the leukemic bone marrow.^27–31^ Our communication results therefore point at the reduced communication within the immune compartment, potentially as a consequence of the immune suppression caused by the tumor.

The top upregulated interactions included interactions between *PDE1B* positive HSPC and three cell types: B-cells, T-cells and monocytes, all expressing the calmodulin gene family (CALM1-3). *PDE1B* positive HSPC have been characterized as leukemia initiating cell populations^32^. Although calmodulin expression was unchanged in the immune cells, the change in communication was driven by the increase in all three components of HSPC: the cell type abundance, the active fraction of HSPC expressing *PDE1B,* as well as the mean expression level of *PDE1B* within the active fraction (Fig. 3e,g; Supplementary Fig 6f).

To independently reproduce our results on a different dataset, we applied the same analysis to an integrated AML and healthy dataset with a higher number of samples33,34 (Supplementary Fig. 8a-e). This cohort consisted of 9 AML and 26 healthy samples, with 6 broad cell types (HSPC, monocytes, erythrocytes, B-cells, T-cells and DCs). Again, we found more downregulated than upregulated interactions in AML, and the downregulated interactions included interactions among the immune cells themselves whereas the upregulated interactions included interactions with HSPCs (Supplementary Fig. 8f-h). Using only the shared cell types between the two datasets, we compared the overlap between the interactions in the immune compartment and found 345 mutually differential interactions (out of 1,587 in Lasry and 3,509 in van Galen-Oetjen) (Supplementary Fig. 9a). The mutually downregulated interactions among immune cell types showed a reduction of immune activating signals in the bone marrow, specifically driven by the drop of the monocytes expressing *NRG1* and *HP* (Supplementary Fig. 9b).

In summary, using *community* we show that in AML there is a general downregulation of communication within the immune compartment indicating immuno-suppression, whereas the HSPC compartment containing the tumor cells shows upregulation of communication.

### Interplay of the individual components during communication change

*Community* interaction strength calculation is purposely built in a modular manner, to allow the exploration of the mode of action that leads to a communication change. Namely, for each interaction, *community* considers three components: the cell type abundance of the sending and the receiving cell types (*ρS*, *ρR*), the active fraction of the sending and the receiving cell types for the specific ligand and receptor (or adhesion molecule pair) (*ϕlS*, *ϕrR*), and the expression level of the sending and the receiving cell types for the ligand and the receptor pair in the active cells (alternatively, the adhesion molecule pair) (*πlS*, *πrR*). For any given interaction, there are several possible scenarios which can lead to differential communication between cases and controls, which can easily be disentangled with *community* (Fig. 4a, Supplemental Fig. 10). First, there is a scenario with no change in any of the six components, in which case we speak of a stable interaction. Second, only one of the six components can change (decrease or increase). If strong enough, this change can lead to a statistically significant change of the whole interaction weight, which we address as a simple decrease or increase. Third, several of the six components can change. This change can either be concordant or discordant. In concordant changes all affected components change in the same direction. Here, if the combined change is strong enough, it can also lead to a significant change in the overall interaction strength. Alternatively, the change in the several components can be discordant, i.e. the components change in opposite directions. In this case we can speak of a compensation scenario. If the overall interaction strength changes significantly, we can interpret it as an insufficient compensation. If the overall interaction strength stays stable, we can regard it as a sufficient compensation (Fig. 4a, Supplemental Fig. 10). For both insufficient and sufficient compensation, if the changes are happening only within the sending or only within the receiving population, we speak of self-compensation. If the changes are happening in both the sending and the receiving population, we speak of peer-compensation.

**Fig. 4.**
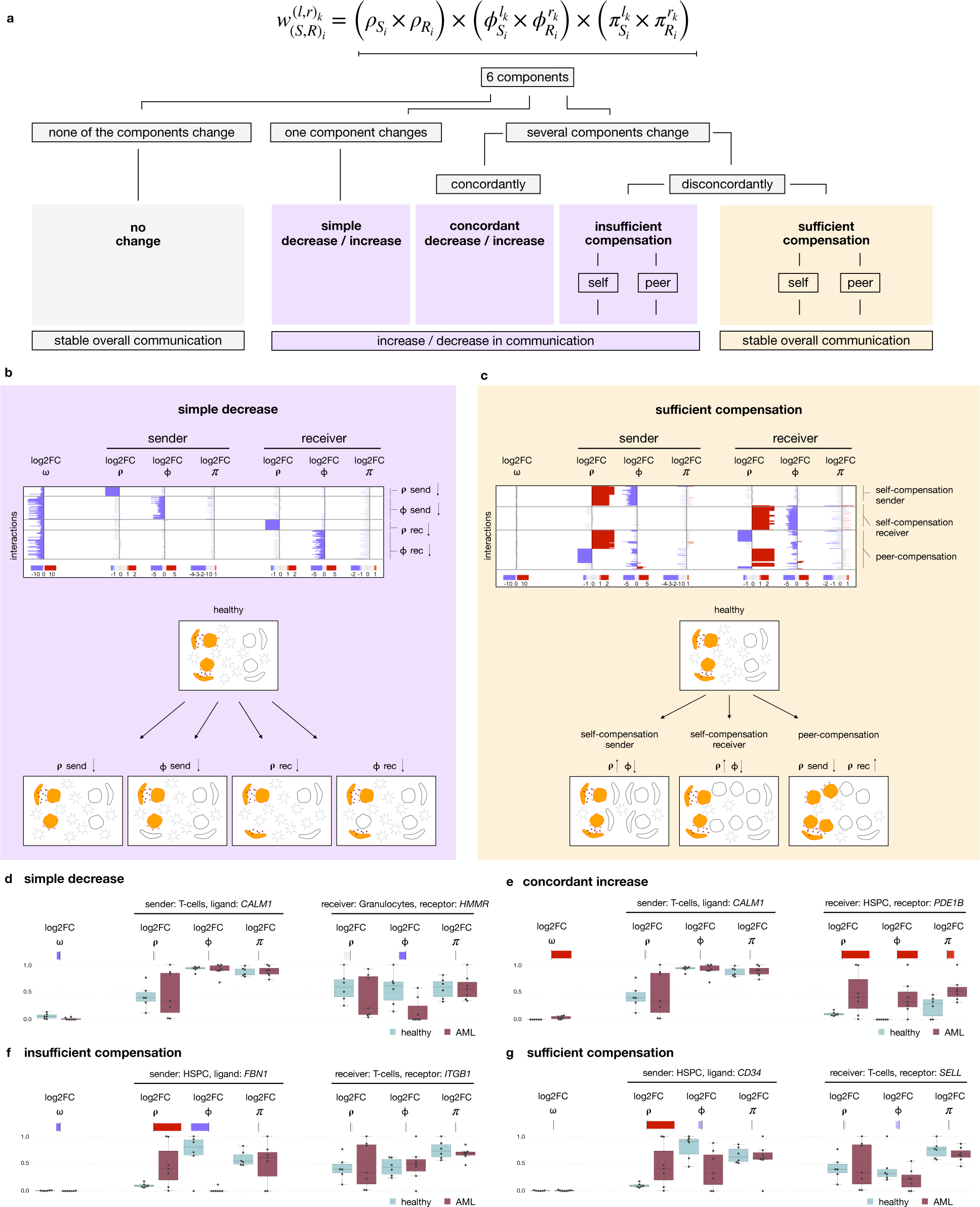
Interplay of individual components shapes the resulting communication shift. **a**, Six types of changes in communication, driven by the interplay of the individual components. **b**, Lasry et al. interactions that fall into the category simple decrease. **c**, Lasry et al. interactions that fall into the category sufficient compensation. **d-g**, Example of an interaction showing simple decrease (T:CAML1_Gran:HMMR) (**d**), concordant increase (T:CALM1_HSPC:PDE1B) (**e**), insufficient compensation (HSPC:FBN1_T:ITGB1) (**f**), and sufficient compensation (HSPC:CD34_T:SELL) (**g**). Upper panels show forest plot for the log2FC of the individual components of the corresponding interactions, color scheme is same as in **b**. Lower panels show box plots with the values for the individual components.

We analyzed the behavior of the individual components in the AML-healthy dataset^23^ and found cases for all categories described above (Fig. 4b, Supplementary Fig. 10). In the dataset, 3405 interactions were truly unchanged (Supplementary Fig. 10). Further 517 interactions were driven by a change in only one component, as in the case of *T:CALM1_Gran:HMMR* where the change was driven by the drop in the active fraction of the receiver (Fig. 4b,d, Supplementary Fig. 10). We observed a concordant change in 1139 interactions, such as in *T:CALM1_HSPC:PDE1B* driven by the increase of all three components of the receiver (Fig. 4e, Supplementary Fig. 10). Further 695 interactions showed discordant changes in the individual components, leading to insufficient compensation, as in the case of *HSPC:FBN1_T:ITGB1*, where despite the increased cell type abundance of the HSPC, the drop of their active fraction expressing FBN1 lead to the decrease on the intensity of this interaction (Fig. 4f; Supplementary Fig. 10). Then, very interestingly, we could identify 1570 instances of compensated interactions. For example, in the interaction *HSPC:CD34_T:SELL* there was an increase of the cell type abundance of the HSPC and a decrease of the active fraction of the HSPC expressing *CD34*, as well as the active fraction of the T cells expressing *SELL*. The overall communication in this interaction was not significantly affected due to the peer-compensation (Fig. 4g; Supplementary Fig. 10).

The ability to distinguish between the situation of true stable communication and the sufficiently compensated communication is an important and unique feature of *community*. Compensated interactions can for example be detectable alterations observed in the early stages of a disease which can potentially get decompensated during the disease progression.

### Method comparison

We compared *community* to two popular cell-cell communication tools: CellPhoneDB^6^ (v3.1.0) and NicheNet^10^ (v2.0). Although initially the tools were developed to study static communication in a sample, both recently provided a pipeline for differential communication analysis which is based on differential gene expression analysis using Seurat^35^ (v4.3.0).

To assess the performance of all tools, we first compared the ligand-receptor pair databases that each of them provides. We unified the database formats by splitting complexes into individual components (see Methods) and enforced the directionality of ligand-receptor pairs, with ligand being the first component of the pair and receptor being the second (Fig. 5a). Both *community* and NicheNet included two levels of confidence: curated, that is cited in the literature, or predicted, that is suggested by protein-protein interaction databases. CellPhoneDB only included curated pairs. We compared the full-mode databases, i.e. curated and predicted together, as well as the curated only mode. We performed the comparison on genes, as well as on pairs. *Community* showed the highest number of genes in both curated only and full modes, as well as the highest number of pairs in curated only mode. NicheNet showed the highest number of pairs in the full mode, this large number was driven by the predicted pairs. *Community* shows the second highest number of pairs in the full mode (Fig. 5b).

**Fig. 5.**
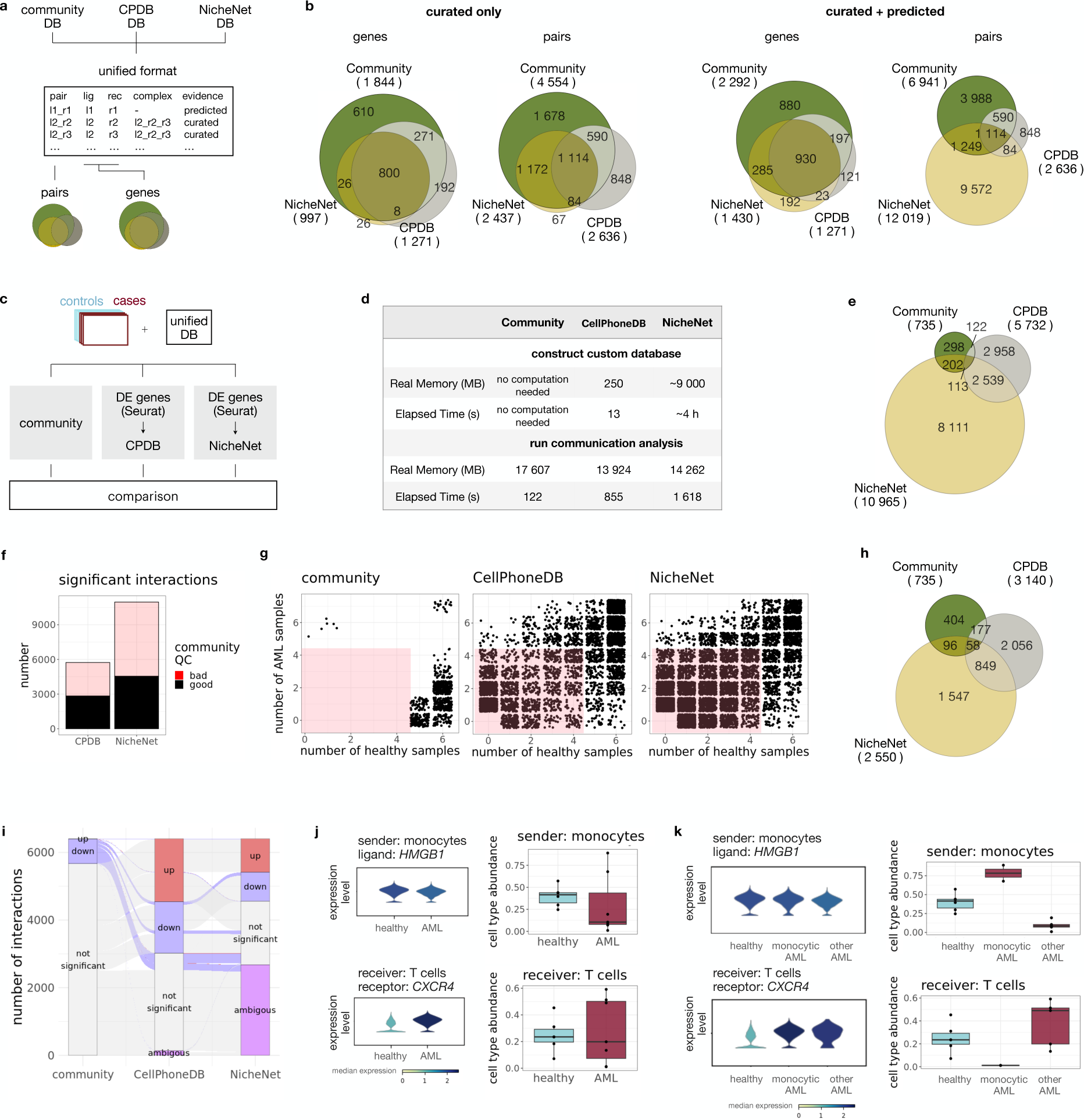
Comparison to existing pipelines. **a**, Pipeline to compare the ligand-receptor databases of *community,* CellPhoneDB, and NicheNet. **b**, Comparison of number of genes (i.e. ligands and receptors) and number of ligand-receptor pairs in the curated only and curated+predicted databases of *community*, CellPhoneDB, and NicheNet. **c**, Pipeline to compare the communication results of *community,* CellPhoneDB, and NicheNet. **d**, Real memory and elapsed time comparison. **e**, Differential interactions. **f**, Differential interactions of CellPhoneDB and NicheNet that were considered as bad or good quality by the *community* QC. **g**, Differential interactions identified by *community*, CellPhoneDB, or NicheNet showing in how many healthy or AML samples an interaction was found (each dot is an interaction). Red area represents the region identified as bad quality (under-represented) by *community*. **h**, Differential interactions after removing the under-represented differential interaction from CellPhoneDB and NicheNet. **i**, Alluvial plot of the union of the interactions in panel **h** stratified by up- or down regulated, ambiguous, and not significant. **j**, An example of an interaction (Mono:*HMGB1* to T:*CXCR4*) exhibiting a stabilising compensatory effect via cell type abundance. Left panels show expression levels of *HMGB1* and *CXCR4* in monocytes and T cells respectively. Right panels show cell types abundance of monocytes and T cells in healthy and AML samples. **k**, Mono:*HMGB1* to T:*CXCR4* interaction when regarding the two monocytic AML cases as a separate subtype. Left panels show expression levels of *HMGB1* and *CXCR4* in monocytes and T cells respectively. Right panels show cell type abundance of monocytes and T cells in healthy and AML split by subtype.

We next compared the differential communication results of *community* to the Seurat-based application of NicheNet and CellPhoneDB using the Lasry et al. dataset^23^ (Fig. 3a). To ensure that the differences in the results are driven by the algorithm behavior, and not by the differences in the original ligand-receptor databases, we used the *community* database for all tools, which we converted to the tool-specific formats (Fig. 5c). As customizing the database, e.g. adding new ligand-receptor pairs, might be important for some studies, we measured the time and memory consumption of this step for CellPhoneDB and NicheNet, using their provided pipelines. The custom database construction took 2 minutes and 5 hours time and 250MB and 9GB memory for CellPhoneDB and NicheNet, respectively (Fig. 5d).

For both CellPhoneDB and NicheNet, we applied the provided differential communication pipelines, starting with the computation of the differentially expressed (DE) genes between the cases and the controls for each cell type separately using Seurat. This resulted in the following numbers of DE genes: 386 for monocytes, 337 for granulocytes, 246 for T-cells, 472 for NK, 297 for B-cells, 452 for HSPC, 1,082 for erythrocytes, and 239 for DCs (Supplementary Table 2). CellPhoneDB and NicheNet differential communication analysis pipelines resulted in 5,732 and 10,965 differential interactions respectively (Fig. 5e). Since *community* applies an additional quality control step before calculating differential interactions (see sections Theoretical Basis and Methods), it gave the most conservative number of 735 differential interactions. Indeed, nearly a half of differential interactions in both CellPhoneDB and NicheNet were also detected by *community,* but they were filtered out as they did not pass the quality control (Fig. 5f). During this quality control step, *community* removes interactions that are present in too few samples (i.e. less than 5 in either case or control cohort). On the contrary, both the CellPhoneDB and NicheNet pipelines apply the thresholds on the minimum number of active cells per cell type and gene on the integrated dataset. This means that in some occasions a gene in an interaction will pass the threshold (for example, 10% of active cells in the cell type, which is the default threshold in both methods) even if it is expressed by cells that only belong to one sample. Since neither CellPhoneDB nor NicheNet implement additional QC filters, such interactions are captured in the final result making it hard to distinguish patient individual changes from disease specific communication changes. To quantify this effect, we calculated the number of samples (healthy vs AML) where an interaction was detected, per method. *Community* only kept the interactions that were found in at least five healthy or AML samples, whereas CellPhoneDB and NicheNet captured many interactions which were present even in just one sample (Fig. 5g). When we removed these under-represented interactions, CellPhoneDB and NicheNet showed only 3,140 and 2,550 differential interactions respectively (Fig. 5h).

While *community* assigns the differential interactions to up- or down-regulated, CellPhoneDB and NicheNet do not provide information on the directionality. To further compare the consistency of the results, we assigned the interactions of CellPhoneDB and NicheNet as up- or down-regulated if either one or both genes within the ligand-receptor pair were differentially up- or down-regulated, respectively. In cases where both partners were differentially expressed, but in discordant directions, the interaction was assigned to the class “ambiguous”. NicheNet also identified some interactions as differential where neither the ligand nor the receptor were among differentially expressed genes. These interactions were also assigned as “ambiguous”. All three algorithms showed concordancy in the assignment of direction for all overlapping interactions (Fig. 5i).

Since CellPhoneDB and NicheNet pipelines rely on the mean expression level of genes within a cell type, but are not taking into consideration the cell type abundance, they leave out potential compensation effects caused by shifts in the cell type composition, which *community* instead is able to capture. An example of such an interaction, which is captured by CellPhoneDB as upregulated, but by *community* as unchanged, is between the monocytes expressing *HMGB1* and the T-cells expressing *CXCR4*. *HMGB1*, together with *CXCL12,* has been shown to facilitate *CXCR4*-mediated recruitment of immune cells^36^. Indeed, the expression of *CXCR4* in the T-cells of the AML cohort was significantly higher than in the healthy cohort, while the expression of *HMGB1* by the monocytes remained unchanged (Fig. 5j). Thus, expression-based analysis, as in CellPhoneDB and NicheNet, indicated the increase of the interaction between the two cell types. Yet, the fraction of both cell types in the AML cohort showed a higher interquartile range of values compared to the healthy samples, indicating potential heterogeneous changes among the AML samples. We indeed identified two patients with an AML of monocytic origin (samples AML-0024 and AML-0160) (Supplementary Fig. 6c). While the fraction of monocytes in the “monocytic” AML subtype was elevated compared to the healthy cohort, the T-cells had a significant drop. In this case, despite the elevated receptor expression on the T-cells, there were overall too few T-cells to increase the *HMGB1-CXCR4*-based interaction with the monocytes (Fig. 5k). The “non-monocytic” AML cases showed an opposite development: the fraction of the monocytes dropped compared to the healthy cohort, while the T-cells were partially increased. In this case, despite the increased expression of the receptor in the T-cells, the lack of ligand expressing monocytes may have a compensatory effect on the strength of this interaction, which was captured by *community*. Overall, *community* was able to capture this development and the compensation effect within the monocyte:*HMGB1*_T-cells:*CXCR4* interaction for the AML cohort, despite the two distinct mechanisms taking place (Fig. 5k).

Finally, we tested the resource usage for all three tools using all three datasets: the UC vs healthy from Smillie et al., and the two AML vs healthy datasets from Lasry et al., and the integrated one from van Galen et al. and Oetjen et al. We subsetted each into differing numbers of samples (n=2 to n=40) and measured the elapsed time, as well as peak VRAM usage. *Community* performed as the fastest algorithm in all testing conditions, and it had the best scalability with the increase in the number of samples. *Community* was also the most memory-efficient tool for the biggest dataset (Fig. 5d, Supplementary Fig. 11a-d).

## Discussion

*Community* is a fast and robust R package for differential communication analysis in large scRNAseq case-control studies. *Community* explores the information on three different aspects of the dataset: the shrinkage or expansion of individual cell types (cell type abundance), the alteration in their active fraction, and the gene expression level. The overall change in communication for a particular pair of cell types using a particular ligand-receptor pair depends on the interplay of these components. Due to this feature, *community* can easily track back which components influenced the communication shift most, and by this gain a better understanding of the underlying biological processes.

Due to this detailed component analysis, *community* is able to distinguish between truly unchanged interactions, where none of the components changed, and compensated interactions, where components change in opposite directions compensating each other and keeping the overall communication intensity stable. These communication-stabilizing effects may play a particularly important role in the early stages of diseases where tissue homeostasis can be restored and preserved by such mechanisms^1^.

The ability of detecting compensated interactions is a completely novel feature that *community* offers. One of the key information that *community* uses in this case is the cell type abundance. Most conventional techniques omit this information, focusing only on the expression level of ligands and receptors within cell types. This causes a misleading assignment of differential interactions in cases of compensated communication. For example, if a sending cell type keeps the ligand expression on a stable level, but becomes more abundant, it will result in an increased overall availability of a ligand in the tissue. The receiver cell type may sense this due to the overstimulation of the receptor-associated downstream pathway and activate a feedback mechanism to decrease the level of the receptor to the level and bring the pathway activity back to the physiological level. The interaction between the sender and the receiver will thus be compensated. In this case, it is crucial to regard the information about the sender cell type abundance increase together with the decrease in receptor expression by the receiver, because otherwise the interaction will be identified as downregulated, instead of compensated. This information may be crucial in studies focused on identifying novel disease targets, since disrupting a mechanism that is compensatory may lead to worsening of the disease.

*Community* is able to detect changes in the active fractions of cell types. This property is especially of advantage in cases where the fraction of cells within a cell type expressing a ligand or receptor is very small. The tools which rely on the differential expression analysis of the ligands and receptors may miss these interactions. This is because the DEG analysis pipeline may not detect these genes, as they are dominated by a large pool of non-expressing cells. For example, in UC, an interaction between the inflammatory monocytes expressing *OSM* and inflammatory fibroblasts expressing *OSMR* has been described37,38. Being specific to a very small subtype of the monocytes and fibroblasts, these genes do not appear among the DEG and thus would be missed by DEG-based techniques. Yet *community* was able to correctly identify this interaction as significantly upregulated.

The information on changes in expression levels within the active fraction of cells serves as an indication of potential shifts in the pathways upstream of the expressed ligands or receptors. This may be due to the altered chromatin accessibility, transcription factor activity or post-transcriptional modifications. *Community* makes it easy to screen for these interactions providing a list of candidates for further analyses with orthogonal techniques.

*Community* provides a large database of curated and predicted ligand-receptor interactions and adhesion molecules. In contrast to other tools, *community* keeps the information about complex molecules in the database, yet splits them for the calculation of the individual interaction. The tools that keep the composition of a complex fixed most often calculate the average expression over the complex components. If an interaction decreased or increased in disease, with this strategy it requires additional analysis to track back whether this change affected all components of a complex or just some of them. In contrast, with *community* keeping track of each component of the complex separately, it is easy to reconstruct the driving changes, as we show with the example of the *Myeloid:OSM_Fibroblasts:OSMR-IL6ST* interaction in the Smillie et al. dataset. Another phenomenon is that the composition of the complex may alter in disease compared to healthy39,40 or among different cell types40,41. In this case, if by chance the average expression value of this complex stays similar, this compositional change will be missed by conventional methods, but captured by *community*.

Another strength of *community* is its robustness against outlier samples. Especially when dealing with human data, heterogeneity and individual-specific signatures is a frequently encountered issue, both due to sample preparation and interindividual differences. The conventional pipelines for DEG analysis in scRNAseq with several samples in cases and controls like Seurat^35^ or scanpy^42^ do not distinguish between cells coming from different samples and are thus sensitive to outlier samples. As a consequence, the communication analysis techniques discover putatively differential communication signatures that are driven by these outlier samples. In contrast, *community* does not merge all the cells from a cohort in one pool, but rather reconstructs communication in each sample separately. This makes it easy to discard outlier sample-driven interactions during the QC and differential communication analysis step, to robustly identify communication changes that are generally driven by the disease.

Collectively, *community* is a user-friendly and scalable R tool to run differential communication analysis in scRNAseq-based case-control type of analyses. *Community* produces robust cell-cell interaction analysis, providing information on potential biological processes underlying the changes in communication in disease, such as expansion or shrinkage of certain cell types, the active fraction producing the ligand or the receptor, and their expression levels within the active cells. *Community* is fast, memory efficient and scalable, and provides intuitive visualization. In summary, *community* is a tool for analyzing large datasets and better understanding cell-cell communication processes in health and disease.

## Methods

### *Community* database

To construct *community* database, two datasets from OmniPath^11^ were used: *ligrec extra*, containing ligand-receptor interactions without literature reference (total of 8350 interactions as of February 2024), *curated ligand-receptor interactions* (total of 5141 interactions as of February 2023). Both binary and complex interactions within the *ligrec extra* and *curated ligand-receptor interactions* datasets were considered. Complex interactions were broken down into binary interaction as follows. First, a complex interaction is broken down into all possible binary pairwise combinations. Next, these pairs were checked against the protein-protein interaction (PPI) network of OmniPath to ensure that the identified binary interactions are supported by existing PPI data. A gene was categorized as a true ligand or receptor, if it was classified as such by at least two resources. Genes that did not fall in the ligand or a receptor category were classified as adhesion molecules. The default order of genes in a ligand-receptor pair was set such that the ligand is first and the receptor is the second. If a ligand-receptor interaction was found in a swapped position, we correct the order accordingly. For adhesion molecule interactions, the consensus directionality among the resources was checked in *ligrec extra* and *curated ligand-receptor interactions*. In case of consensus agreement, the consensus directionality was retained, otherwise, the directionality followed lexicographical order. Additionally, functional annotation for each gene was also added to the dataframe using the MyGene library (v1.38.0; Mark A. et al., 2023), providing users with additional information. The final database included 6941 pairs.

### Interaction weight formula

A weight of an interaction is calculated for a particular sample and a particular ligand-receptor (or adhesion molecule) pair between two cell types. The weight *ω* of an interaction for a sample between the sending cell type expressing a ligand of a pair *k* receiving cell type expressing a receptor from the pair *k* is calculated as:

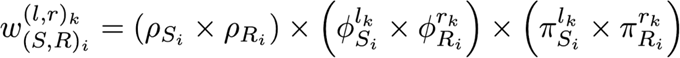

where *ρ_Si_* and *ρ_Ri_* are the relative cell type abundance of the sending cell type (*ρ_Si_*) and the receiving cell type (*ρ_Ri_*); *ϕ^lk^_Si_* and *ϕ^rk^_Ri_* are relative active fraction of the ligand in the sending cell type and the receptor in the receiving cell type; *π^lk^_Si_* and *π^rk^_Ri_* are relative mean expression of the ligand in the active fraction of the sending cell type and the receptor in the active fraction of the receiving cell type. See supplemental methods for details.

### *Community* pipeline

To calculate communication, three thresholds were set: threshold_celltype_size, threshold_nr_active_cells, and threshold_expr. The threshold_celltype_size is a threshold for the minimum number of cells in a cell type in one sample. If the number of cells in the cell type of interest in a particular sample is less or equal to the threshold_celltype_size, this cell type is considered missing in this sample. The threshold_nr_active_cells is a threshold for the minimum number of active cells in a cell type in one sample. An active cell is defined as a cell expressing a gene over the threshold_expr (see below). If the number of active cells for a specific gene in a cell type in one sample is smaller or equal to the threshold_nr_active_cells, then the active fraction of this cell type for this gene in this sample is considered to be zero. The threshold_expr is a threshold for an expression value of a gene in a single cell. If an expression value of a gene in a cell is greater than threshold_expr, this cell is considered active. Otherwise, the expression of this gene in this cell is set to zero and the cell is considered inactive for this gene.

For the quality check, the following filters are used: the cumulative interaction weight filter, the presence per cohort filter and the ligand/receptor expression filter. An interaction is considered of good quality, if it passes all three filters. The cumulative interaction weight filter checks the log10 cumulative weight over all samples computed for each interaction and removes the low weight interactions. To pass this filter, the interaction needs to be greater than the threshold_log10_cum_weight threshold. This threshold is set at the elbow of probability distribution density of the log10 cumulative weights. The presence per cohort filter checks the fraction of samples in which an interaction was detected (i.e. has a non-zero value) in the control cohort and, separately, in the case cohort and removes interactions found in too few samples. To pass this filter, an interaction needs to have a greater value than the threshold_frac_samples_per_condition threshold either in the control cohort or in the case cohort or in both. The ligand/receptor expression filter checks the mean expression level of the ligand and the receptor of an interaction in the case and the control samples (separately) and removes the interactions with lowest expressed ligands or receptors or both. This filter uses a threshold_log10meanexpr_per_condition threshold. For each interaction, the following four log10 mean expression values are checked: i) the ligand in sending cells in control samples, ii) the receptor in receiving cells in control samples, iii) the ligand in sending cells in case samples, and iv) the receptor in receiving cells in case samples. An interaction passes this filter if both its ligand and receptor pass the threshold either in control samples or in case samples or in both.

The differential communication is calculated on good quality interactions between the case and the control cohort using either t-test^12^ on log2-transformed weights or Wilcoxon test^13^ on non-transformed weights. The default adjustment method for multiple testing is FDR. The log2FC of an interaction is calculated as log2 of the ratio of its mean weight in the cases over its mean weight in the controls. To identify significantly differential interactions, the and the should be specified by the user.

### Preprocessing of healthy - ulcerative colitis dataset

The dataset consists of 48 biopsies taken from the colon of 12 healthy and 18 ulcerative colitis (UC) individuals^15^, several biopsies were taken per individual, yet to exclude co-dependencies in the data, only one biopsy was kept per individual (see below) (accession number SCP259). For the UC individuals, only inflamed biopsies were considered. Epithelial and lamina propria layers were combined for each sample. The original 50 cell types were merged to 19 broader categories. (Suppl. Table 1) During quality control, only cells with 1,000 to 30,000 reads and more than 500 genes were kept. For gene filtering, a pseudo bulk count threshold greater than 3 was set. Each cell type needed to have a minimum of 5 cells in each sample and be captured in at least 40 out of 48 samples. The sample filter was set to 12 out of 13 cell types. For each individual, only the biopsy having the highest number of cell types and cells was kept. This yielded 13,861 genes, as well as 160,482 cells, 13 cell types, and 11 healthy and 17 UC samples. The filtered data was normalized by cell type using scran^43^ (v1.26.2). Batch correction was performed using *scGen^44^* (v2.1.0).

### Run *community* on healthy - ulcerative colitis dataset

*Community* was run on log-transformed normalized counts. The threshold_celltype_size and threshold_nr_active_cells were both set to >6, and threshold_expr >0.1. The total number of interactions were 491,959. For the quality control the threshold_log10_cum_weight was selected to be >0.03, the threshold_frac_samples_per_condition >0.5 and threshold_log10meanexpr_per_condition >0.015. The total number of good quality interactions was 15,675. Finally, the threshold_log2FC was set to >1 and threshold_fdr was set to <0.1, yielding a total of 758 downregulated and 2,612 upregulated interactions.

### Preprocessing of healthy and AML bone marrow dataset

ScRNAseq communication analysis of the bone marrow of healthy and AML individuals was performed on publicly available data^23^. The log-transformed normalized read counts and the annotation files were downloaded from GSE185381. In the AML cohort, only samples at initial diagnosis were kept. Duplicated samples from healthy individuals #4 and #5 were removed. This yielded 14 samples. In the first pre-processing step, we removed genes that had all-zero values, yielding 31,843 genes. Cell IDs that were not present in the cell annotation files were excluded, yielding 58,354 cells.

In the quality control step, filters on cells, cell types, genes, and samples were applied. To exclude bad quality cells, the library size per cell was set to be between 1,100 and 30,000 reads and the number of genes was set to greater than 500. To filter cell types, first original cell subtypes were pulled into 11 bigger classes (Suppl. Table 1). A good quality cell type was defined as having at least 5 cells in each sample and being captured in at least 12 samples. Megakaryocytes, perivascular cells and lymphoid progenitor (lymP) cells types did not pass these thresholds and were excluded. Genes having a cumulative pseudobulk count less than 0.25 were excluded, yielding 15,770 genes. Samples having had less than 7 cell types were excluded, yielding 13 samples, 8 cell types and 46,702 total cells.

### Run *community* on healthy and AML bone marrow dataset

*Community* was run on log-transformed normalized counts. For the communication analysis, the threshold_celltype_size and the threshold_nr_active_cells were both set to 6 cells, the threshold_expr was set to 0.1. The total number of interactions were 151,744. For the QC, the threshold_log10_cum_weight, the threshold_frac_samples_per_condition, and the threshold_log10meanexpr_per_condition were set to 0.01, 0.6, and 0.02 respectively. The total number of good quality interactions was 22,645. The differential communication analysis was run on good quality interactions using t-test on log2-transformed weights and FDR adjustment. The threshold_log2FC was set to 1 and the threshold_fdr was set to 0.1, which yielded 2,555 downregulated and 109 upregulated interactions.

### Preprocessing of integrated datasets from two human AML-healthy bone marrow studies

An integrated dataset containing the scRNAseq of the bone marrow of healthy individuals (GSE116256^33^, GSE120221^34^) and AML patients at initial diagnosis (GSE116256^33^) was created. From the GSE120221 dataset, the samples S1, Sk1, S2, Ck, C2 were removed as they duplicates. From the dataset GSE116256, the BM5-34p, MUTZ3, and OCI-AML3 samples, as well as samples with less than 50% blasts were removed. In the first pre-processing step, the genes that were missing in one of the datasets, as well as a cumulative pseudobulk count less than 1 were removed. Cell IDs that were not found in the cell annotation files were excluded. The library size per cell was set to be between 1,000 and 30,000 reads and the number of genes was set to be greater than 500. Additionally, cells that formed a cluster of high library size and low gene number area were removed by applying a linear threshold of 3,000log^10^(library size + 1) - 10,500. Original cell types were pulled into 8 bigger classes (Suppl Table 1). The mutation bearing cells (marked as “-like”) and their healthy counterparts were merged into the same cell type categories. A well represented cell type was defined as the one having at least 5 cells in each sample and being captured in at least 30 samples. Samples which had less than 5 cell types were excluded. This yielded 12,485 genes, 74,583 total cells, 6 cell types and 33 samples. The data was normalized using *scran^43^* (v1.30.2) on each cell type separately, preserving natural cell type differences. The batch effect using *scGen^44^* (v2.1.0).

### Run *community* on two integrated healthy and AML bone marrow datasets

Community was run on log-transformed normalized and batch corrected counts. For the communication analysis, the and the threshold_nr_active_cells were both set to 6 cells, and the threshold_expr was set to 0.1. The total number of interactions were 62,064. For the QC, the threshold_log10_cum_weight, threshold_frac_samples_per_ condition, and the threshold_log10meanexpr_per_condition were set to 0.01, 0.6, and 0.02 respectively. The total number of good quality interactions was 7,043. The differential communication analysis was run on good quality interactions using t-test on log2-transformed weights with FDR adjustment. The threshold_log2FC was set to 1 and the threshold_fdr was set to 0.1, which yielded 3,280 downregulated and 229 upregulated interactions.

### Method comparison for cell-cell interaction tools

To ensure that the results of different algorithms were not driven by the differences in the databases, the comparison was run on the *community* database, which was brought to the algorithm-specific format. For CellPhoneDB^6^, three main CSV files were built: *gene_input*, *protein_input*, and *interactions*. For NicheNet^10^, three individual networks were built: *ligand-receptor interactions* was built using *community* database, *signaling interactions* was built using all the protein-protein interactions datasets from OmniPath while removing self-interactions, *gene regulatory network* was built using DoRothEA^45^ regulons available through OmniPath, with confidence levels ranging from A to C. The optimization was done via mlrMBO (v1.1.5.1) as described in the NicheNet workflow.

Method comparison was performed on the log-transformed normalized counts of the *Lasry et. al.* dataset. To identify differentially expressed genes (DEGs) per cell type between cases and controls, the FindMarkers module (min.pct=0.10) from Seurat^35^ library was run on each cell type separately (v4.3.0). To count as DEG, the average log-2-fold change(avg_log2FC) has to be greater than 0.25 or less than -0.25 and the adjusted p-value has to be < 0.05 (Supplemental Table 2).

CellPhoneDB was run on each sample separately with method *degs_analysis* flag along with the list of differentially expressed genes (DEGs) sourced from Seurat and a *--threshold* of cells expressing the ligand or receptor set to 0.10. The directionality of the output pairs was double checked and put according to the original database, if swapped. NicheNet was run on each individual sample and each possible pair of cell types serving as sender and receiver. An additional expression threshold of a gene being expressed in at least 10% of cells of a cell type in a sample was used, `get_expressed_genes` function was utilized with *pct* parameter set to 0.10. DEGs for the receiver cell type were passed as a gene set of interest. The potential ligands, which are the ligands expressed by the sending cell type and have corresponding receptors in the receiving cell type, were pooled and then used as input to NicheNet. The 30 top-ranked ligands were used for downstream analysis. The receptors of the top-ranked ligands were obtained from the *optimized weighted network of ligand-receptor interactions*. Finally, we combine each cell type from each sample and unify the structure, where rows are corresponding to an interaction while columns are samples, and values are corresponding scores. *Community* was run with the following thresholds: the threshold_celltype_size and the threshold_nr_active_cells were both set to 6 cells, and the threshold_expr was set to 0.1. For the QC, the threshold_log10_cum_weight, the threshold_frac_samples_per_condition, and the threshold_log10meanexpr_per_condition were set to 0.01, 0.6, and 0.02 respectively. As for the differential communication analysis, threshold_log2FC was set to 1 and the threshold_fdr was set to 0.1. An additional threshold of genes being expressed in at least 10% of a cell type in a sample was used to mimic the CellPhoneDB and NicheNet pipelines.

To measure the computational time and memory usage, “psrecord” (v1.3) was used. All algorithms were run on one computational node of the server, equipped with 36 Cores, 256 GB RAM and running CentOS Linux 7 (Core).

## Code availability

*Community* package and the tutorials can be downloaded from https://github.com/SoloveyMaria/community. The data preprocessing pipelines can be found at https://github.com/colomemaria/community-paper.

## Data availability

The raw and the pre-processed datasets from Lasry et al. are available at https://www.ncbi.nlm.nih.gov/geo/query/acc.cgi?acc=GSE185381 and https://zenodo.org/records/7962808 respectively. The raw datasets from van Galen et al. and Oetjen et al. are available at https://www.ncbi.nlm.nih.gov/geo/query/acc.cgi?acc=GSE116256 and https://www.ncbi.nlm.nih.gov/geo/query/acc.cgi?acc=GSE120221. The processed integrated dataset is available at https://zenodo.org/records/10013368. The raw dataset from Smillie et al. can be accessed at Broad DUOS, a controlled-access data repository, https://singlecell.broadinstitute.org/single_cell/study/SCP259/intra-and-inter-cellular-rewiring-of-the-human-colon-during-ulcerative-colitis, while the pre-processed version available at https://zenodo.org/records/10013294

## Author Contributions

M.S., M.C.T, F.Z., and A.S. designed the study. M.S, M.A.C., F.R.S., M.I., and M.A.-F. performed the package development and data analysis. M.S, M.C.T., F.Z., M.A.C, and F.R.S interpreted the results. M.S, M.C.T, and F.Z wrote the manuscript with input from all authors.

## Supporting information

Suppplemental Table 2

## Acknowledgements

We acknowledge the Core Facility Statistical Consulting of Helmholtz Zentrum Munich, and in particular Dr. Marina Jimenez Munoz, Ronan Le Gleut, Raphael Meixner, and Prof. Dr. Christian Fuchs, for the consulting on the statistical testing. The present contribution is supported by the Helmholtz Association under the joint research school “HIDSS-006 - Munich School for Data Science@Helmholtz, TUM&LMU. This work was supported by the Impuls-und Vernetzungsfonds of the Helmholtz-Gemeinschaft (grant VH-NG-1219) for M.C.T. and M. S. F.Z. acknowledges funding from the Förderprogramm für Forschung und Lehre (grant FöFoLe-1048) of the LMU University as well as from the German Cancer Consortium (DKTK) and the Bavarian Cancer Research Center (BZKF).

---

**Supplemental Figure 1.**
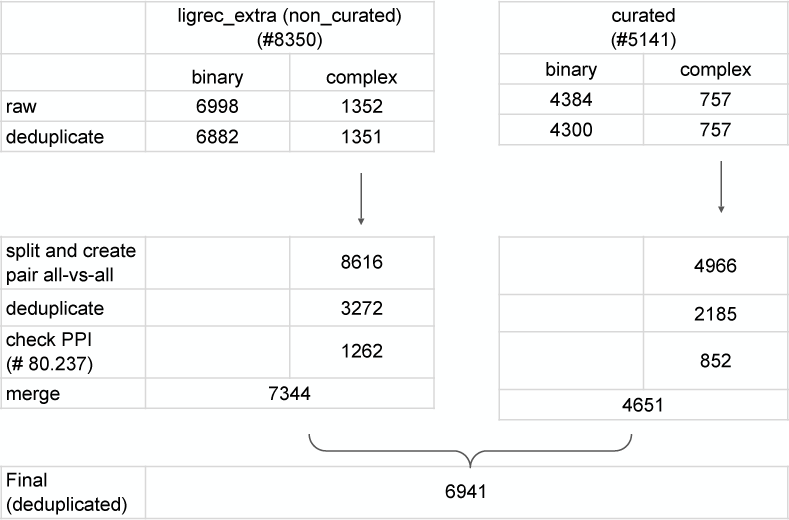
Community ligand-receptor database.

**Supplemental Figure 2.**
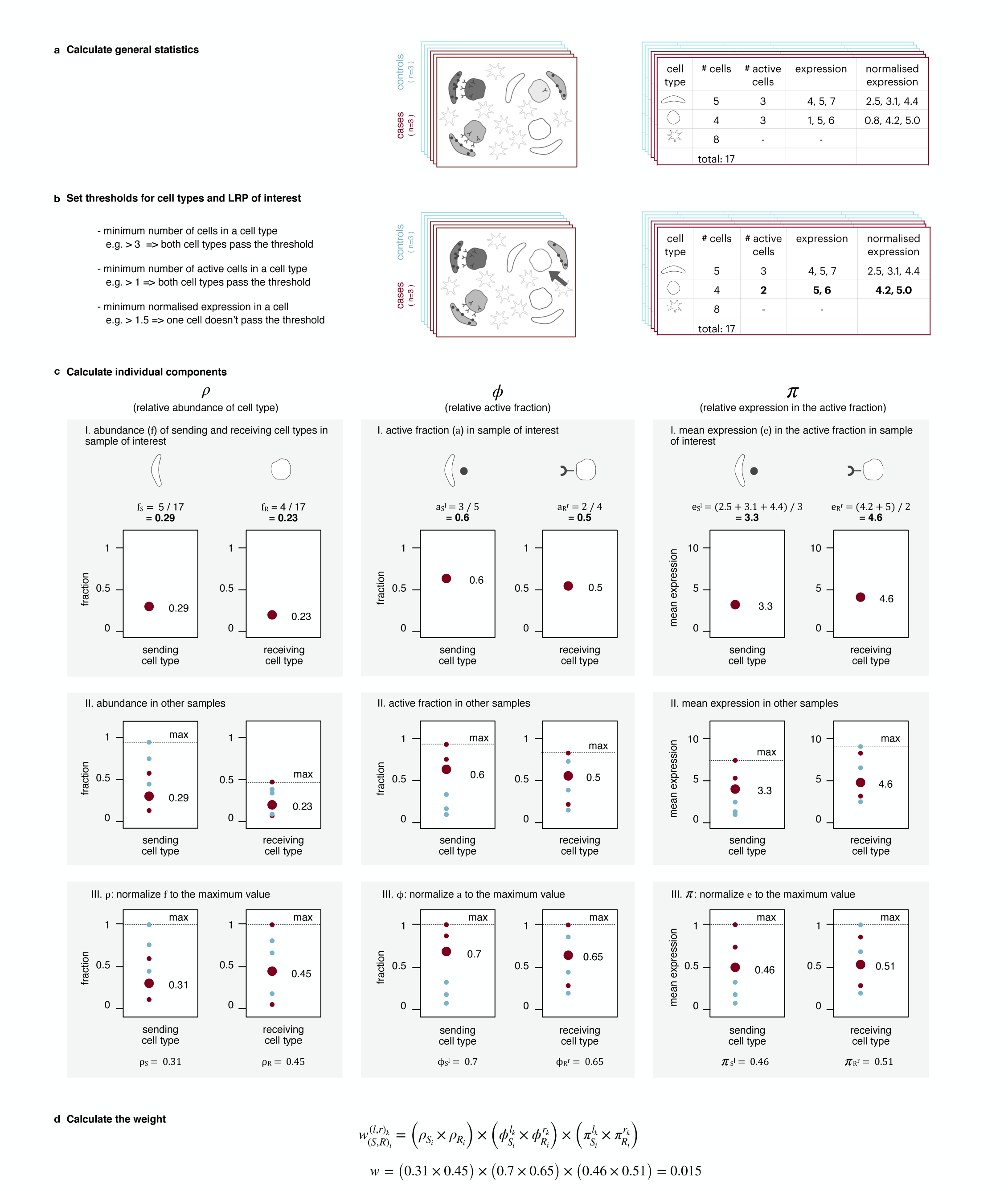
Formula to calculate individual interaction weight between a sending cell type of interest expressing a ligand of interest and a receiving cell type of interest expressing a receptor of interest. **a**, Step 1: calculate general statistics for each cell type in each sample: number of cells, number of active cells (i.e. cell expressing the ligand or receptor of interest respectively). **b**, Step 2: set and apply thresholds: i) minimum number of cells in a cell type, ii) minimum number of active cells in a cell type, iii) minimum normalised expression in a cell. **c**, Step 3: calculate the relative cell type abundance, relative active fraction, and relative mean expression in the active fraction for the sending and the receiving cell types. **d**, Step 4: multiply all six value to get the interaction weight.

**Supplemental Figure 3.**
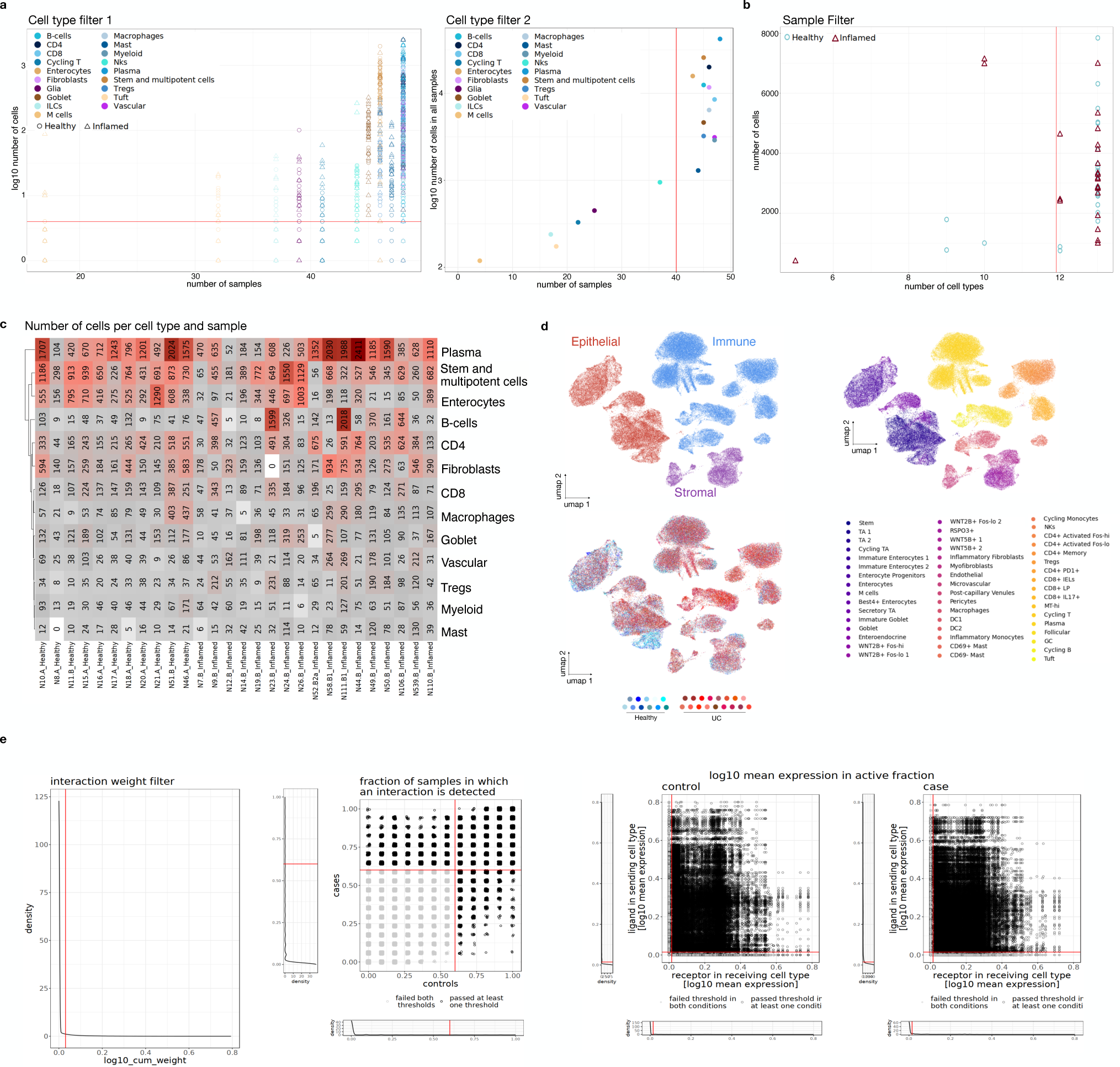
Quality check in the Smillie et al. dataset. **a**, Cell type filters. The red line represents the filtering thresholds: each cell type in each sample should contain at least 5 cells per cell type (left panel) and each cell type must be present in at least 40 samples (right panel). **b**, Sample filter. Each sample should contain at least 12 cell types (red line presents the threshold). **c**, Number of cells per cell type and sample after filtering. **d**, UMAP visualization of the dataset after batch correction colored by tissue type (top left), original cell types (top right) and samples (bottom left). **e**, QC of the interactions: interaction weight filter (far left panel), fraction of samples in which an iteration is detected (middle left panel), mean expression of the ligands and the receptors in the active fraction of the corresponding cell types in the control cohort (middle right panel) and the case cohort (far right panel).

**Supplemental Figure 4.**
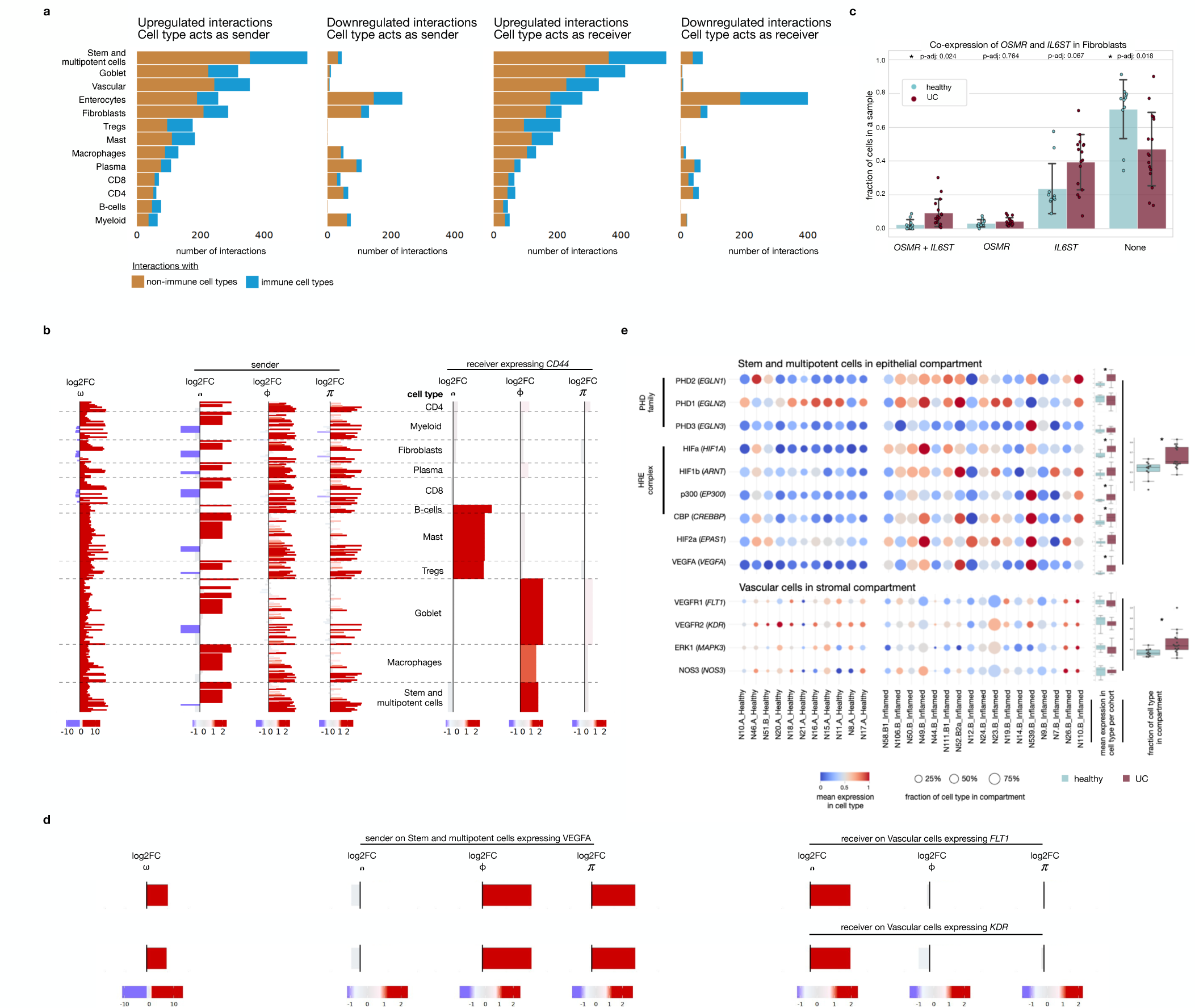
Interaction analysis of Smillie et al. data. **a**, Number of significant (p-val < 0.05) interactions per cell type, split depending on if the cell type acts as a sender, or a receiver and if the interactions are up- or downregulated. The colore indicates whether the counterpart cell types belong to the non-immune compartment (gold) or to the immune compartment (blue). **b**, Forest plot showcasing the individual components of significantly up- and downregulated interactions with *CD44* as a receiver. **c**, Bar plots showcase the distribution of the genes OSMR and IL6ST over all Fibroblasts. P-values were calculated using t-tests (p-val. <0.05), with an asterisk marking significance. The error bars were calculated with the python function sem() for the standard error of the mean. Bonferroni correction was performed. **d**, Forest plot showing the individual interactions between VEGFA on Stem and multipotent cells (left side) and FLT1/KDR on Vascular cells(right side), split by components. **e**, Dot-plot of *VEGFA, EPAS1, HRE complex and PHD family* expression in stem and multipoint cells and *FLT1, KDR, MAP3K and NOS3* expression in vascular cells. The color represents mean gene expression for a cell type per sample. Dot-size represents the fraction of the respective cell type in the cell-specific compartment. Statistics were done by performing t-tests. Significant findings (p-val. <0.05) are labeled with an asterisk.

**Supplemental Figure 5.**
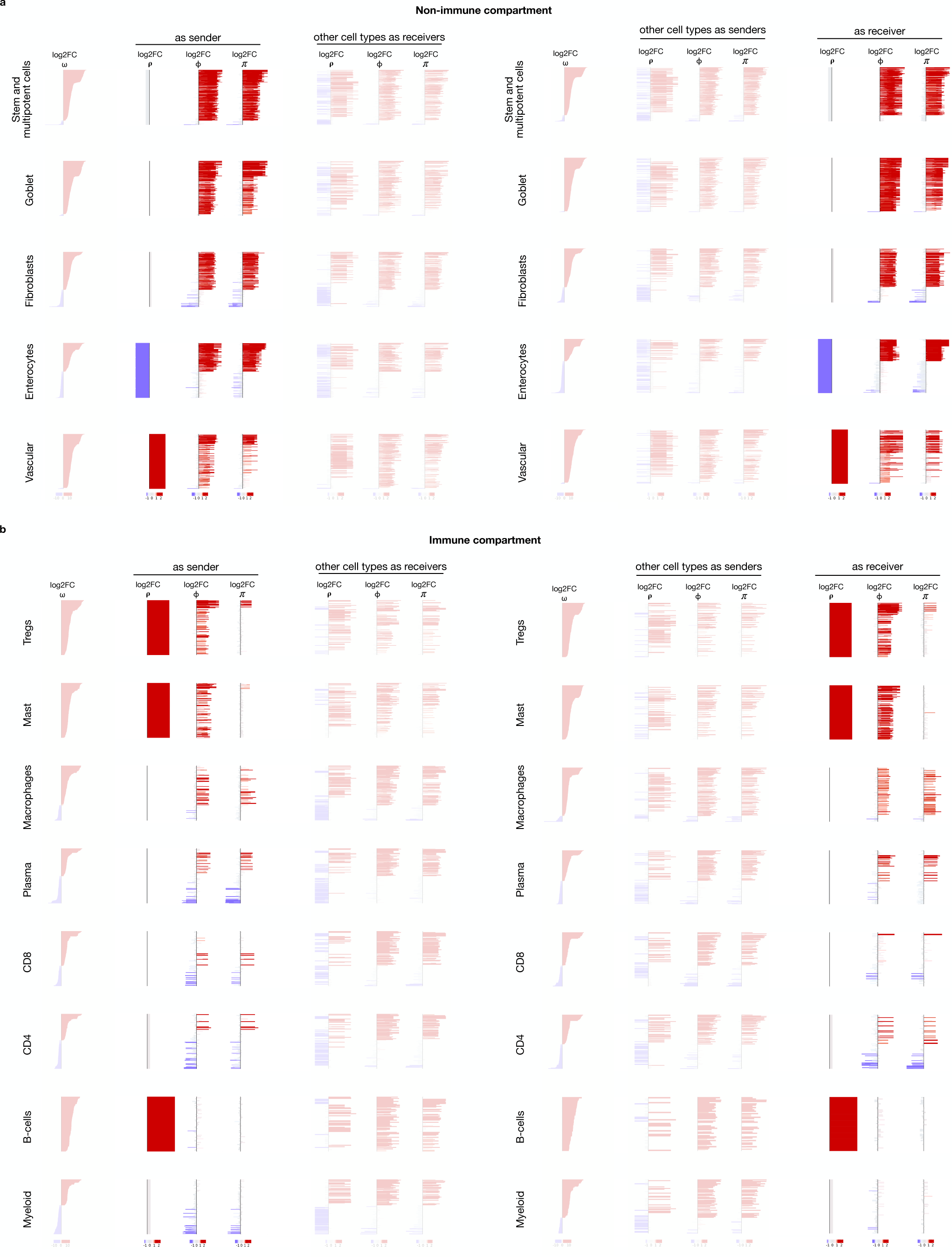
Differential interactions in Smillie et al. data split by components and cell types. The figure shows all significantly up- and downregulated interactions for the non-immune (**a)** and immune (**b)** cell types. For each cell type, two forest plots are plotted: a plot where the cell type acts as a sender (left part) or as a receiver (right part). Each forest plot contains log2 fold change representation for the individual components ρ, ϕ and *π*, as well as the total interaction weight *w*.

**Supplemental Figure 6.**
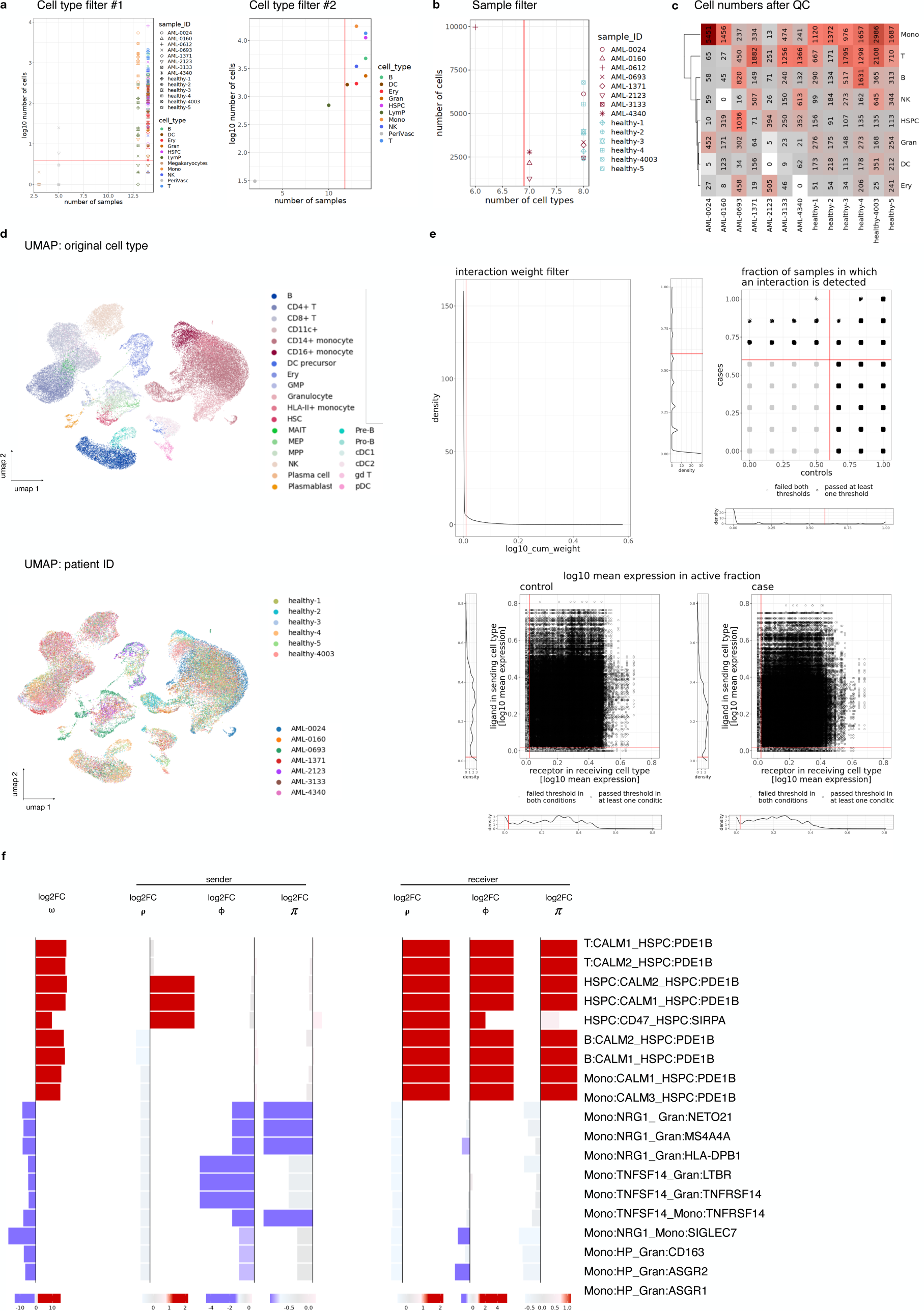
Quality check in the Lasry et al. dataset. **a**, Cell type filters. The red line represents the filtering thresholds: each each cell type in each sample should contain at least 5 cells per cell type (left panel) and each cell type must be present in at least 12 samples (right panel). **b**, Sample filter. Each sample should contain at least 7 cell types (red line presents the threshold) **c**, Number of cells per cell type and sample after filtering. **d**, UMAP coloured by the original cell types annotation from the Lasry et al. publication (upper panel) and patient ID (lower panel). **e**, QC of the interactions: interaction weight filter (upper left panel), fraction of samples in which an iteration is detected (upper right panel), mean expression of the ligands and the receptors in the active fraction of the corresponding cell types in the control cohort (lower left panel) and the case cohort (lower right panel). **f**, Forest plot showcasing the individual components of top up-and down-regulated interactions.

**Supplemental Figure 7.**
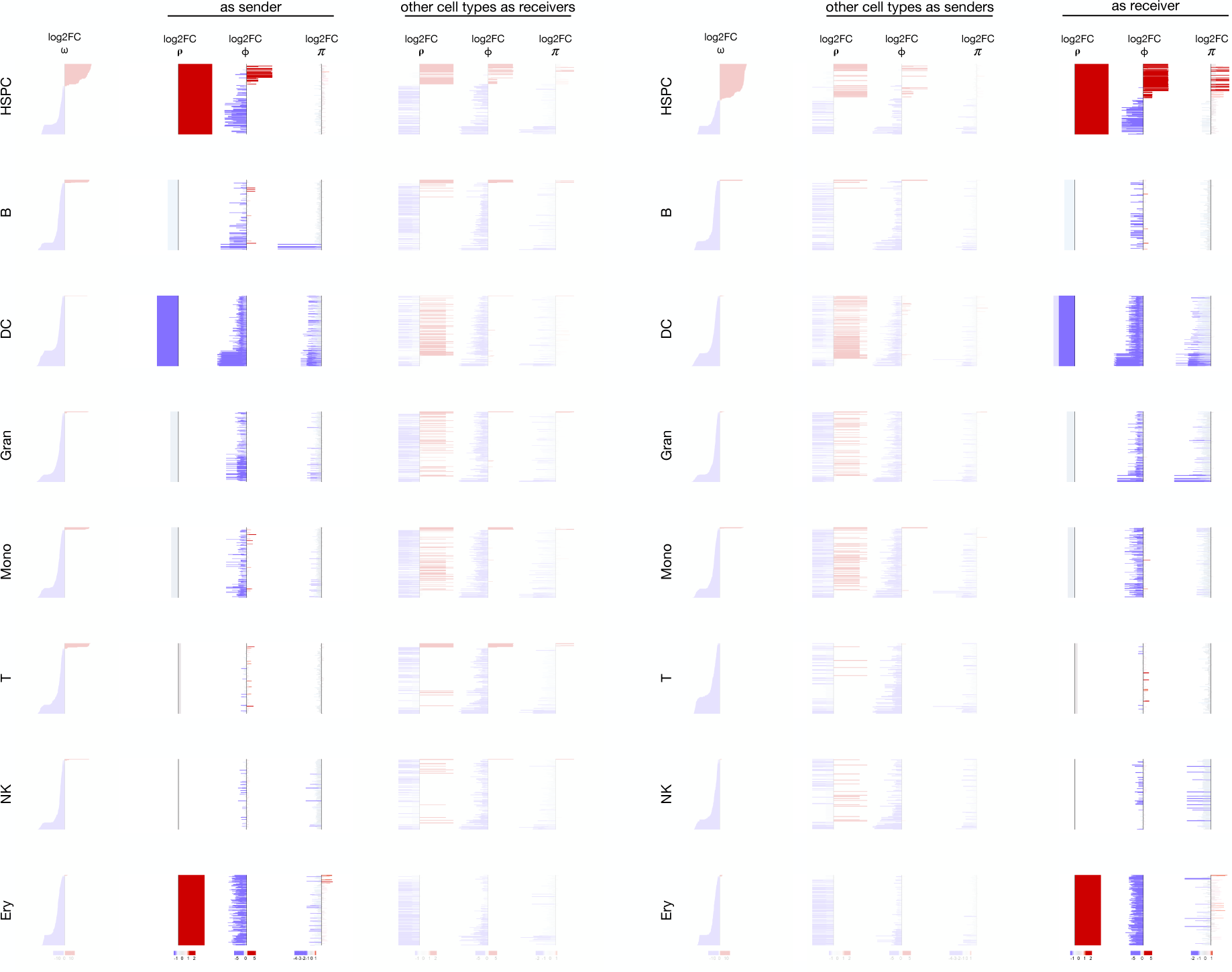
Interactions in Lasry et al. data split by components and cell types. The figure shows all significantly up- and downregulated interactions for all cell types. For each cell type, two forest plots are plotted: a plot where the cell type acts as a sender (left part) or as a receiver (right part). Each forest plot contains log2 fold change representation for the individual components ρ, ϕ and *π*, as well as the total interaction weight *w*.

**Supplemental Figure 8.**
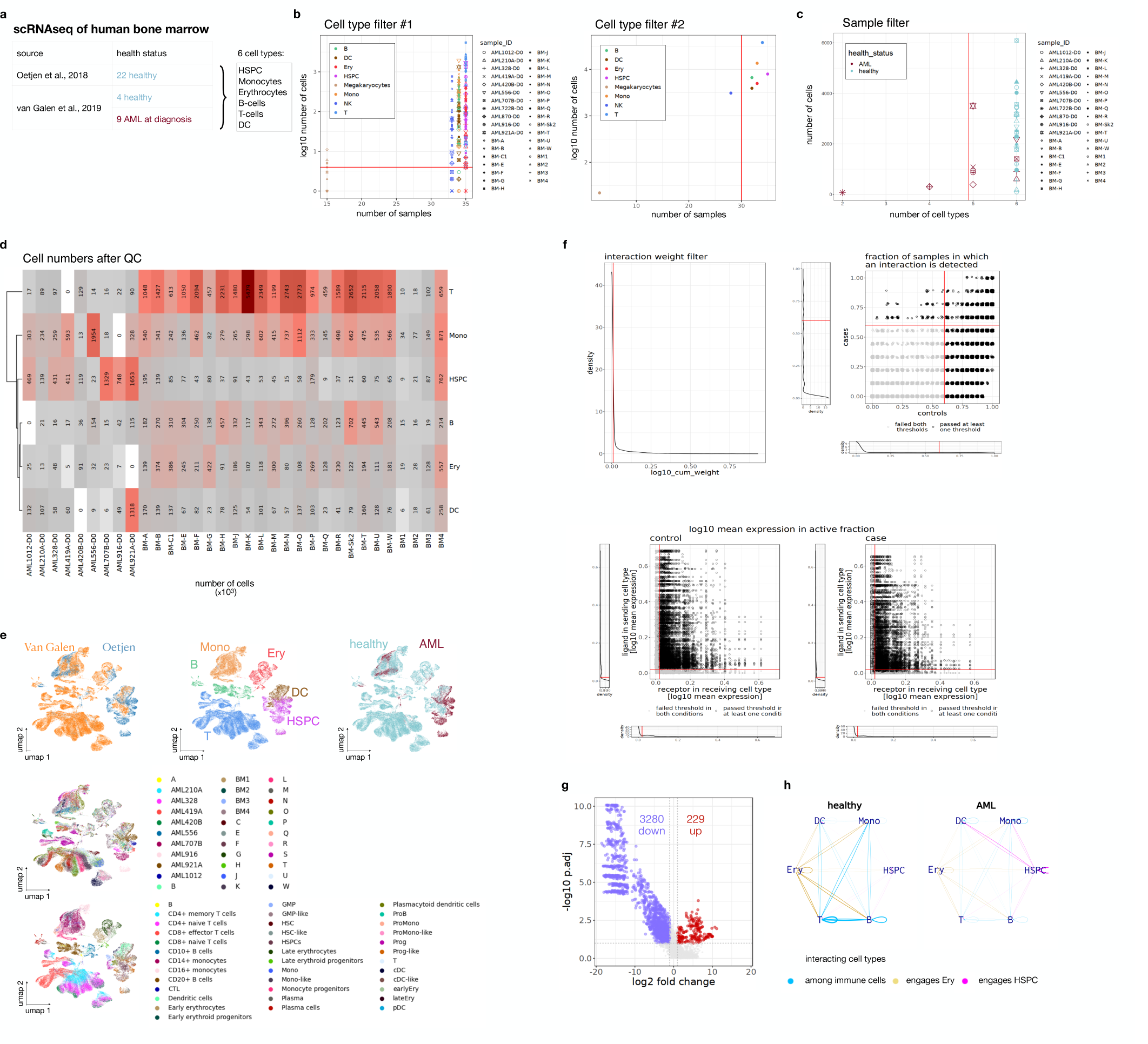
Quality check in the integrated van Galen et al. - Oetjen et al. dataset. **a**, Overview of both scRNAseq datasets used for human bone marrow analysis. The table summarizes the source of the datasets, health status of the samples, and the 6 merged cell types. **b**, Cell type filters. The red line represents the filtering thresholds: each cell type in each sample should contain at least 5 cells per cell type (left panel) and each cell type must be present in at least 30 samples (right panel). **c**, Sample filter. Each sample should contain at least 5 cell types (red line presents the threshold). **d**, Number of cells per cell type and sample after filtering. **e**, UMAP visualization of the dataset after batch correction colored by dataset (top left panel), cell type (top center panel), health status (top right panel), samples (middle panel) and original cell types (bottom panel). **f**, QC of the interactions: interaction weight filter (upper left panel), fraction of samples in which an iteration is detected (upper right panel), mean expression of the ligands and the receptors in the active fraction of the corresponding cell types in the control cohort (lower left panel) and the case cohort (lower right panel). **g**, Volcano plot of differential interactions (AML vs healthy). **h**, Network graph of differential interactions of healthy (left panel) and AML (right panel) cohort. One line represents cumulative weight of all differential interactions between the two connected cell types.

**Supplemental Figure 9.**
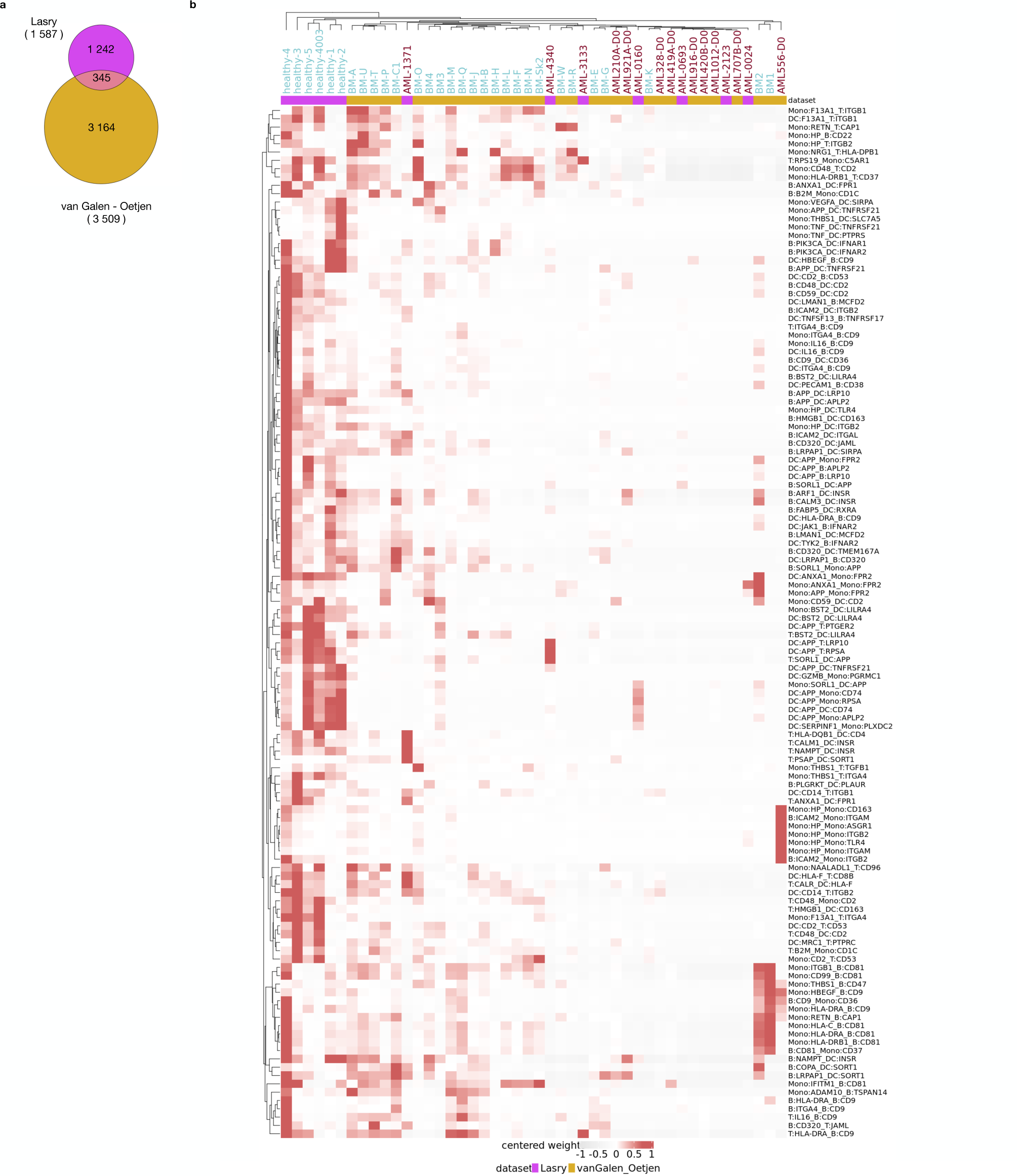
Overlap between differential interactions of the Lasry (granulocytes and NKs excluded) and the integrated van Galen et al. - Oetjen et al. datasets. **a**, Euler diagram of the differential interactions of Lasry et al and integrated van Galen et al. - Oetjen et al. **b**, Heatmap of the 124 overlapping differential interactions among the immune cell types.

**Supplemental Figure 10.**
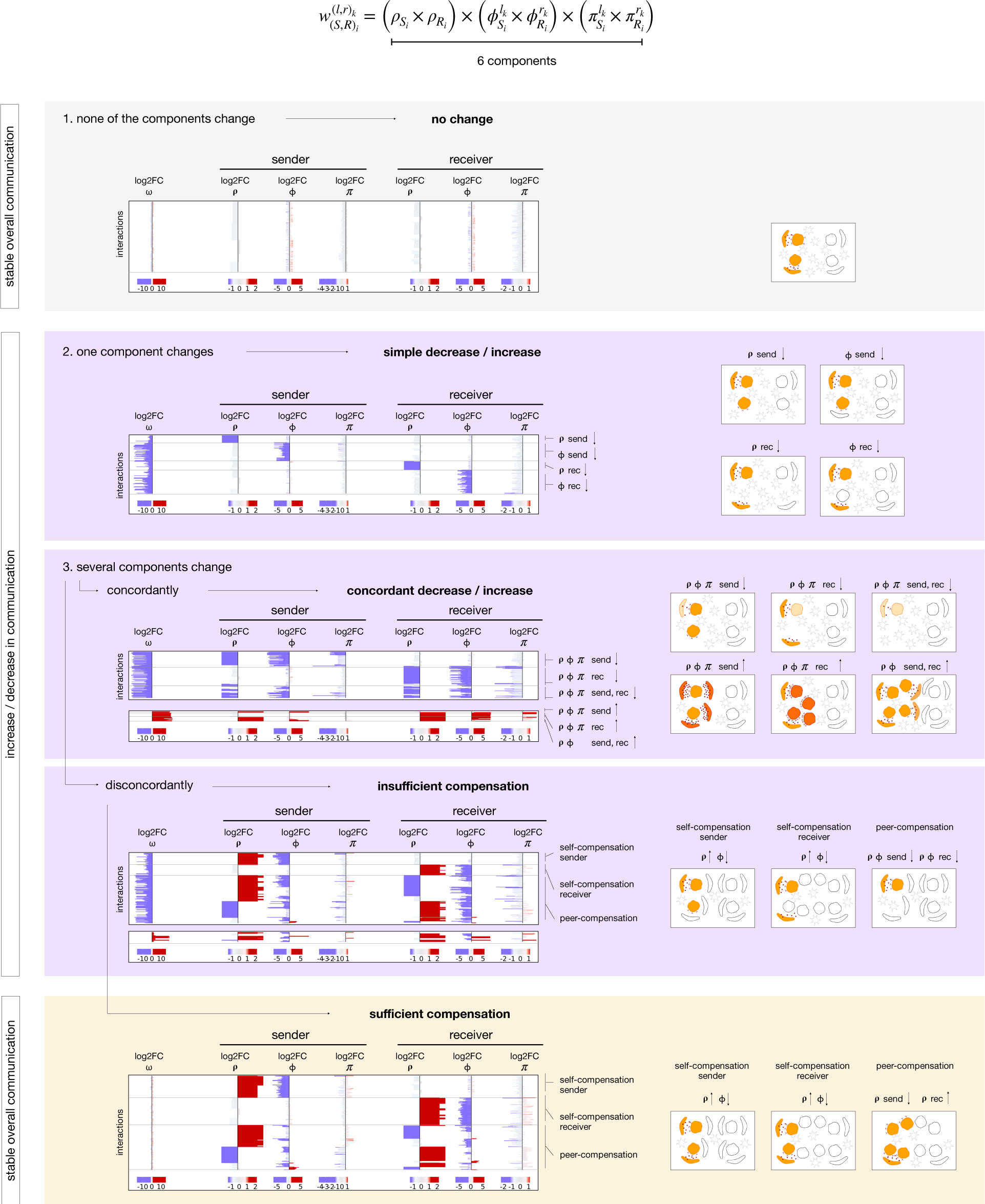
Interplay of individual components shapes the resulting communication shift.

**Supplemental Figure 11.**
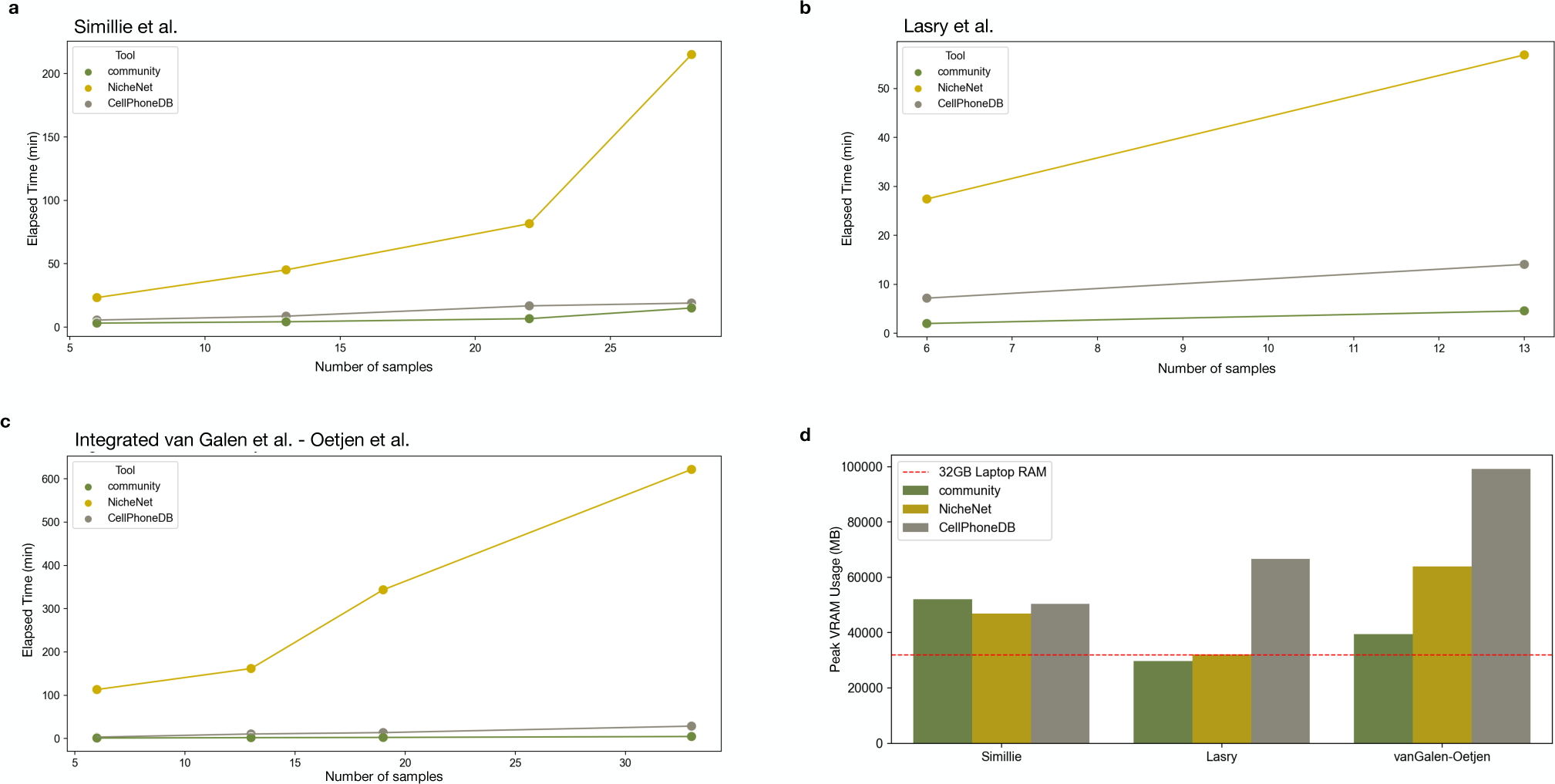
Resource usage. a-c,Benchmarking of elapsed time for the *community*, NicheNet and CellPhoneDB tools across the three datasets with with increasing number of samples: Smillie et al. (a), Larry et al. (b), and integrated van Galen et al. - Oetjen et al. (c) d, Comparison of peak RAM usage among the tools and across three full datasets: 22 samples for Simillie et al., 13 samples for Lasry et al., and 33 samples for the integrated van Galen-Oetjen datasets. The dashed red line indicates the capacity of 32 GB, a specification often seen in laptops used for data-intensive tasks.

## References

1. Armingol, E., Officer, A., Harismendy, O. & Lewis, N. E. Deciphering cell-cell interactions and communication from gene expression. Nat. Rev. Genet. 22, 71–88 (2021).

2. Bechtel, T. J., Reyes-Robles, T., Fadeyi, O. O. & Oslund, R. C. Strategies for monitoring cell-cell interactions. Nat. Chem. Biol. 17, 641–652 (2021).

3. Tanay, A. & Regev, A. Scaling single-cell genomics from phenomenology to mechanism. Nature 541, 331–338 (2017).

4. Valls, P. O. & Esposito, A. Signalling dynamics, cell decisions, and homeostatic control in health and disease. Curr. Opin. Cell Biol. 75, 102066 (2022).

5. Solovey, M. & Scialdone, A. COMUNET: a tool to explore and visualize intercellular communication. Bioinformatics (2020) doi:10.1093/bioinformatics/btaa482.

6. Efremova, M., Vento-Tormo, M., Teichmann, S. A. & Vento-Tormo, R. CellPhoneDB: inferring cell-cell communication from combined expression of multi-subunit ligand-receptor complexes. Nat. Protoc. 15, 1484–1506 (2020).

7. Raredon, M. S. B. et al. Computation and visualization of cell-cell signaling topologies in single-cell systems data using Connectome. Sci. Rep. 12, 4187 (2022).

8. Wilk, A. J., Shalek, A. K., Holmes, S. & Blish, C. A. Comparative analysis of cell-cell communication at single-cell resolution. Nat. Biotechnol. (2023) doi:10.1038/ s41587-023-01782-z.

9. Browaeys, R., et al. MultiNicheNet: a flexible framework for differential cell-cell communication analysis from multi-sample multi-condition single-cell transcriptomics data. bioRxiv 2023.06.13.544751 (2023) doi:10.1101/2023.06.13.544751.

10. Browaeys, R., Saelens, W. & Saeys, Y. NicheNet: modeling intercellular communication by linking ligands to target genes. Nat. Methods 17, 159–162 (2020).

11. Türei, D., Korcsmáros, T. & Saez-Rodriguez, J. OmniPath: guidelines and gateway for literature-curated signaling pathway resources. Nat. Methods 13, 966–967 (2016).

12. Student. The Probable Error of a Mean. Biometrika 6, 1–25 (1908).

13. Wilcoxon, F. Individual Comparisons by Ranking Methods. Biometrics Bulletin 1, 80–83 (1945).

14. Benjamini, Y. & Hochberg, Y. Controlling the false discovery rate: A practical and powerful approach to multiple testing. J. R. Stat. Soc. 57, 289–300 (1995).

15. Smillie, C. S. et al. Intra- and Inter-cellular Rewiring of the Human Colon during Ulcerative Colitis. Cell 178, 714–730.e22 (2019).

16. Alkim, C., Alkim, H., Koksal, A. R., Boga, S. & Sen, I. Angiogenesis in Inflammatory Bowel Disease. Int. J. Inflam. 2015, 970890 (2015).

17. Blander, J. M. Death in the intestinal epithelium-basic biology and implications for inflammatory bowel disease. FEBS J. 283, 2720–2730 (2016).

18. Tanaka, T., Oyama, T., Sugie, S. & Shimizu, M. Different Susceptibilities between Apoe- and Ldlr-Deficient Mice to Inflammation-Associated Colorectal Carcinogenesis. Int. J. Mol. Sci. 17, (2016).

19. Franić, I. et al. Expression of CD44 in Leukocyte Subpopulations in Patients with Inflammatory Bowel Diseases. Diagnostics (Basel) 12, (2022).

20. Dotan, I. et al. The role of integrins in the pathogenesis of inflammatory bowel disease: Approved and investigational anti-integrin therapies. Med. Res. Rev. 40, 245–262 (2020).

21. Hjortebjerg, R., Thomsen, K. L., Agnholt, J. & Frystyk, J. The IGF system in patients with inflammatory bowel disease treated with prednisolone or infliximab: potential role of the stanniocalcin-2 / PAPP-A / IGFBP-4 axis. BMC Gastroenterol. 19, 83 (2019).

22. Richards, C. D. The enigmatic cytokine oncostatin m and roles in disease. ISRN Inflamm 2013, 512103 (2013).

23. Lasry, A. et al. An inflammatory state remodels the immune microenvironment and improves risk stratification in acute myeloid leukemia. Nat Cancer (2022) doi:10.1038/ s43018-022-00480-0.

24. Stubbins, R. J. & Karsan, A. Differentiation therapy for myeloid malignancies: beyond cytotoxicity. Blood Cancer J. 11, 193 (2021).

25. Tettamanti, S., Pievani, A., Biondi, A., Dotti, G. & Serafini, M. Catch me if you can: how AML and its niche escape immunotherapy. Leukemia 36, 13–22 (2022).

26. Hanahan, D. Hallmarks of Cancer: New Dimensions. Cancer Discov. 12, 31–46 (2022).

27. Skeate, J. G. et al. TNFSF14: LIGHTing the Way for Effective Cancer Immunotherapy. Front. Immunol. 11, 922 (2020).

28. Berrocal-Rubio, M. Á. et al. Discovery of NRG1-VII: a novel myeloid-derived class of NRG1 isoforms. bioRxiv 2023.02.02.525781 (2024) doi:10.1101/2023.02.02.525781.

29. Varchetta, S. et al. Engagement of Siglec-7 receptor induces a pro-inflammatory response selectively in monocytes. PLoS One 7, e45821 (2012).

30. Huntoon, K. M. et al. The acute phase protein haptoglobin regulates host immunity. J. Leukoc. Biol. 84, 170–181 (2008).

31. Godoy-Tena, G. et al. Epigenetic and transcriptomic reprogramming in monocytes of severe COVID-19 patients reflects alterations in myeloid differentiation and the influence of inflammatory cytokines. Genome Med. 14, 134 (2022).

32. Lerner, A. & Epstein, P. M. Cyclic nucleotide phosphodiesterases as targets for treatment of haematological malignancies. Biochem. J 393, 21–41 (2006).

33. van Galen, P. et al. Single-Cell RNA-Seq Reveals AML Hierarchies Relevant to Disease Progression and Immunity. Cell 176, 1265–1281.e24 (2019).

34. Oetjen, K. A., et al. Human bone marrow assessment by single-cell RNA sequencing, mass cytometry, and flow cytometry. JCI Insight 3, (2018).

35. Hao, Y. et al. Integrated analysis of multimodal single-cell data. Cell 184, 3573– 3587.e29 (2021).

36. Haque, N. et al. Role of the CXCR4-SDF1-HMGB1 pathway in the directional migration of cells and regeneration of affected organs. World J. Stem Cells 12, 938–951 (2020).

37. Liu, J. Z. et al. Association analyses identify 38 susceptibility loci for inflammatory bowel disease and highlight shared genetic risk across populations. Nat. Genet. 47, 979–986 (2015).

38. West, N. R. et al. Oncostatin M drives intestinal inflammation and predicts response to tumor necrosis factor-neutralizing therapy in patients with inflammatory bowel disease. Nat. Med. 23, 579–589 (2017).

39. Egbenya, D. L. et al. Changes in synaptic AMPA receptor concentration and composition in chronic temporal lobe epilepsy. Mol. Cell. Neurosci. 92, 93–103 (2018).

40. Belfiore, A. et al. Insulin Receptor Isoforms in Physiology and Disease: An Updated View. Endocr. Rev. 38, 379–431 (2017).

41. Moruzzi, N. et al. Tissue-specific expression of insulin receptor isoforms in obesity/type 2 diabetes mouse models. J. Cell. Mol. Med. 25, 4800–4813 (2021).

42. Wolf, F. A., Angerer, P. & Theis, F. J. SCANPY: large-scale single-cell gene expression data analysis. Genome Biol. 19, 15 (2018).

43. Lun, A. T. L., McCarthy, D. J. & Marioni, J. C. A step-by-step workflow for low-level analysis of single-cell RNA-seq data with Bioconductor. F1000Res. 5, 2122 (2016).

44. Lotfollahi, M., Wolf, F. A. & Theis, F. J. scGen predicts single-cell perturbation responses. Nat. Methods 16, 715–721 (2019).

45. Garcia-Alonso, L. et al. Transcription Factor Activities Enhance Markers of Drug Sensitivity in Cancer. Cancer Res. 78, 769–780 (2018).

