## supplemental methods for "*Community* assesses differential cell communication using large multi-sample case-control scRNAseq datasets"

### Interaction weight formula

A weight of an interaction is calculated for a particular sample and a particular ligand-receptor (or adhesion molecule) pair between two cell types. We define a ligand as a molecule which is expressed by a sending cell type, and transmits a signal to the receiving cell type. A receptor is expressed by a receiving cell type. A ligand-receptor pair in the database contains can contain only one ligand and one receptor. In the cases where either the ligand or the receptor form a complex, this complex is split into a set of separate one-to-one pairs. The adhesion molecules are treated in the same way as the ligand-receptor pairs. In contrast to the ligand-receptor interaction, we do not assume that an interaction over adhesion molecules has a directionality of the transmitted signal. Yet in the visualization, it might appear as if the directionality were present. In these cases it can be ignored. For simplicity, we refer to both ligand-receptor pairs and adhesion molecule pairs as ligand-receptor pairs.

In the dataset, we have $N$ samples. An in dividual sample is annotated with an index $i \in [1, N]$. A sample $i$ contains $n_i$ total number of cells. The number of cells in the complete dataset is thus

$$\sum_{i=1}^{N} n_i$$

The ligand-receptor pair database contains $K$ ligand-receptor pairs. An individual ligand-receptor pair is annotated with an index $k \in [1, K]$. A ligand of a pair $k$ is annotated as $l_k$ and a receptor as $r_k$.

The dataset contain $C$ different cell types. Ideally, each sample contains the full pallet of $C$ cell types. If a cell type $c_i$ is missing in a sample $i$, all interactions with this cell type will have a zero weight in this sample. Per interaction, we define two cell types of interest: a sending cell type $S$ and a receiving cell type $R$. While calculating communication in a sample $i$, we regard interactions between all possible pairs of cell types (including self-loops), such that each individual cell type $c_i$ acts at least once as a sending cell type $S$ and at least once as a receiving cell type $R$.

In a sample $i$, the number of cells that belong to the sending cell type $S$ is $n_{S_i}$ and the number of cells that belong to the receiving cell type $R$ is $n_{R_i}$. In the same sample, the number of cell of the sending cell type that express a ligand $l$ from a pair $k$ is $n_{S_i}^{l_k}$ and the number of cells of the receiving cell type that express the receptor $r$ from the pair $k$ is $n_{R_i}^{r_k}$.

The weight $w$ of an interaction for a sample $i$ between the sending cell type $S$ expressing a ligand $l$ of a pair $k$ and a receiving cell type $R$ expressing a receptor $r$ from the pair $k$ is calculated as:

$$w_{(S,R)_i}^{(l,r)_k} = (\rho_{S_i} \times \rho_{R_i}) \times \left(\phi_{S_i}^{l_k} \times \phi_{R_i}^{r_k}\right) \times \left(\pi_{S_i}^{l_k} \times \pi_{R_i}^{r_k}\right)$$

The **relative cell type abundance** is defined over the $\rho$ parameter as follows:

$$\rho_{S_i} = \frac{f_{S_i}}{\max\limits_{N(i)}\left\{f_S\right\}}$$

and

$$\rho_{R_i} = \frac{f_{R_i}}{\max\limits_{N(i)}\left\{f_R\right\}}$$

where $f_{S_i}$ and $f_{R_i}$ are the fraction of the sending cell type (or receiving, respectively) in a sample $i$, calculated as

$$f_{S_i} = \frac{n_{S_i}}{n_i}$$

and

$$f_{R_i} = \frac{n_{R_i}}{n_i}$$

The number of cells in a cell type $n_{S_i}$ and $n_{R_i}$ is controlled by the minimum number of cells per cell type threshold. The default value is 4. If $n_{S_i}$ (or $n_{R_i}$) is less than the threshold, then it is set to zero.

The $\max\limits_{N(i)}\{f_S\}$ and the $\max\limits_{N(i)}\{f_R\}$ is the maximum fraction of this sending cell type (or receiving cell type, respectively) over all samples in the dataset. See Supplemental Fig. 1 for schematic visualization.

The **relative active fraction** of cells within a cell type of interest is defined over the $\phi$ parameter as follows:

$$\phi_{S_i}^{l_k} = \frac{a_{S_i}^{l_k}}{\max\limits_{N(i)}\left\{a_S^{l_k}\right\}}$$

$$\phi_{R_i}^{r_k} = \frac{a_{R_i}^{r_k}}{\max\limits_{N(i)}\left\{a_R^{r_k}\right\}}$$

where $a_{S_i}^{l_k}$ and $a_{R_i}^{r_k}$ are the active fraction of the sending cell type $S$ expressing ligand $l_k$ (or receiving cell tpy e $R$ expressing receptor $r_k$, respectively) in a sample $i$, calculated as

$$a_{S_i}^{l_k} = \frac{n_{S_i}^{l_k}}{n_{S_i}}$$

and

$$a_{R_i}^{r_k} = \frac{n_{R_i}^{r_k}}{n_{R_i}}$$

The number of active cells in a cell type $n_{S_i}^{l_k}$ and $n_{R_i}^{r_k}$ is controlled by the minimum number of active cells per cell type threshold. The default value is 0. If $n_{S_i}^{l_k}$ (or $n_{R_i}^{r_k}$) is less than the threshold, then it is set to zero.

The $\max\limits_{N(i)}\{a_S^{l_k}\}$ and the $\max\limits_{N(i)}\{a_R^{r_k}\}$ is the maximum active fraction of this sending cell type expressing ligand $l_k$ (or receiving cell type expressing receptor $r_k$, respectively) over all samples in the dataset. See Supplemental Fig. 1 for schematic visualization.

The **mean expression** $\langle e \rangle$ in the active fraction $\phi$ of the cells is defined over the $\pi$ parameter as follows:

$$\pi_{S_i}^{l_k} = \frac{\langle e_{S_i,\phi}^{l_k} \rangle}{\max\limits_{N(i)}\left\{\langle e_{S,\phi}^{l_k} \rangle\right\}}$$

and

$$\pi_{R_i}^{r_k} = \frac{\langle e_{R_i,\phi}^{r_k} \rangle}{\max\limits_{N(i)}\left\{\langle e_{R,\phi}^{r_k} \rangle\right\}}$$

where $\langle e_{S_i,\phi}^{l_k} \rangle$ and $\langle e_{R_i,\phi}^{r_k} \rangle$ are the mean expression of the ligand $l_k$ in the active fraction $\phi$ of the sending cell type $S$ (or of the receptor $r_k$ in the active fraction of the receiving cell type $R$, respectively) in a sample $i$. The expression in an individual cell in controlled by the threshold for the minimum expression level of a ligand or receptor in a cell. The defauls value of this threshold is 0.05. If a cell does not pass this threshold, its expression is set to zero and it is considered inactive for the ligand $l_k$ (or receptor $r_k$).

The $\max\limits_{N(i)}\left\{\langle e_{S,\phi}^{l_k} \rangle\right\}$ and the $\max\limits_{N(i)}\left\{\langle e_{R,\phi}^{r_k} \rangle\right\}$ is the maximum mean expression of the ligand $l_k$ in the active fraction of this sending cell type (or of the receptor $r_k$ in the active fraction of the receiving cell type, respectively) over all samples in the dataset. See Supplemental Fig. 1 for schematic visualization.
